# A systematic study of HIF1A cofactors in hypoxic cancer cells

**DOI:** 10.1101/2022.08.09.503416

**Authors:** Yuxiang Zhang, Saidi Wang, Haiyan Hu, Xiaoman Li

**Affiliations:** Trinity Preparatory School, Winter Park, Florida, United States of America; Department of Computer Science, University of Central Florida, Orlando, Florida, United States of America; Genomics and Bioinformatics Cluster, University of Central Florida, Orlando, Florida, United States of America; Burnett School of Biomedical Science, College of Medicine, University of Central Florida, Orlando, Florida, United States of America

**Keywords:** HIF1A, hypoxia, motif module, cofactors, TF-TF interaction

## Abstract

Hypoxia inducible factor 1 alpha (HIF1A) is a transcription factor (TF) that forms highly structural and functional protein-protein interactions with other TFs to promote gene expression in hypoxic cancer cells. Despite the importance of these TF-TF interactions, we still lack a comprehensive view of many of the TF cofactors involved and how they cooperate in hypoxic cancer cells. In this study, we systematically studied HIF1A cofactors in eight cancer cell lines and discovered 201 potential HIF1A cofactors. These cofactors were statistically and biologically significant, with 30 of the top 37 cofactors validated or supported in the literature. Moreover, these predicted cofactors include 21 of the 29 known HIF1A TF cofactors in public databases. These discovered cofactors can be essential to HIF1A’s regulatory functions and may lead to the discovery of new therapeutic targets in cancer treatment.

## Introduction

Many human cells require oxygen to carry out their metabolic functions. When those cells sense their oxygen supply is limited, hypoxia occurs ^1^. Cancer cells are a great example of hypoxia, as their ability to rapidly replicate and proliferate forces them to quickly exceed their oxygen supply ^2^. To survive, cancer cells, mainly solid tumors, have developed various defense mechanisms to adapt to hypoxic environments ^3^. Such defense mechanisms primarily rely on the activation of hypoxia inducible factor 1 alpha (HIF1A), a transcription factor (TF) that interacts with other cofactor TFs to activate transcriptional responses like increased glucose uptake and angiogenesis. Since the responses HIF1A regulates facilitate a cancer cell’s survival in hypoxic conditions ^4^, many resources have been diverted in HIF1A targeted treatments ^5,6^.

Though many attempts to target HIF1A have been proposed, one of the most appealing prospects has been to focus on targeting HIF1A through its protein-protein interactions (PPIs). This method has demonstrated high diagnostic potential ^7^. Like many TFs, HIF1A interacts with a multitude of cofactor TFs to effectively control gene expression ^8^. For instance, HIF1A is known to dimerize with its binding partner aryl hydrocarbon receptor nuclear translocator (ARNT) to cooperatively co-regulate target genes. PPIs between different TFs such as HIF1A and ARNT are extremely important due to their ability to confer high specificity and drive differential gene expression ^9,10^. Despite their importance, we found that in many databases containing experimentally curated PPIs – BioGRID ^11^, HPRD ^12^, and BIND ^13^ – very few of the HIF1A cofactors listed were TFs. The undiscovered TF cofactors could have novel therapeutic responses and contribute to a much deeper understanding of hypoxia’s regulatory mechanisms.

In this study, we systematically investigated the potential TF interactions of HIF1A. Measuring possible PPIs in an experimental setup, usually done using co-immunoprecipitation arrays, is time-consuming and costly. Instead, using computational methods to find such cofactors possesses many benefits due to their high recall rate for predicting cofactors in a time-efficient manner ^14–16^. We computationally identified 201 potential HIF1A TF cofactors, many of which were supported by PPI databases and literature. These cofactors were conserved across multiple cell lines and crucial to regulating HIF1A regulated pathways. Our study facilitates an increased understanding of HIF1A’s regulatory functions with transcriptional cooperativity.

## Material and Methods

### Retrieving genomic sequences that HIF1A binds

To systematically study HIF1A cofactors in cancer cell lines, we retrieved raw ChIP-seq data in the following eight human cell lines: 501A ^17^, MDA-MB-231 ^18^, HepG2 ^19^, HKC8 ^19^, HUVEC ^20^, K562 ^21^, LNCaP ^22^, and PC3 ^23^, from the corresponding BioProjects at National Center for Biotechnology Information (NCBI) (Supplementary Table S1). Next, we applied the tool Trimmomatic ^24^ to the raw ChIP-seq read data to trim adaptor sequences and filter low-quality reads. We then mapped the processed reads in each cell line onto the human genome hg38 using Bowtie2 ^25^ and defined the ChIP-seq peaks of the HIF1A binding regions using MACS2 ^26^. To discover TF binding sites (TFBSs) in these ChIP-seq experiments, we extended each MACS2 peak region so that each peak region was at least 800 base pairs long, which is about the median length of a cis-regulatory region ^27,28^. Finally, we extracted the repeat-masked genomic sequences from the UCSC genome browser ^29^ for each peak region. These sequences are likely to contain TFBSs of HIF1A and its cofactors.

### Identifying potential HIF1A TF cofactor motifs and cofactors

To find the potential TF cofactors of HIF1A in a cell line, we applied the tool SIOMICS to the above sequences from the extended ChIP-seq peak regions in this cell line to identify motifs (Fig. 1) ^16^. A motif is a pattern of the TFBSs a TF binds, often represented by a position weight matrix. Many computational tools are developed for motif discovery in ChIP-seq data ^16,30–35^. We chose SIOMICS here because it de novo identifies motifs by discovering motif modules, which can effectively work with large sequence datasets and significantly reduces false positive predictions ^15,16,36^. A motif module is a group of motifs that significantly co-occurs in the input sequences, mimicking the group of motifs for a TF and its cofactors under a given experimental condition. In biological terms, these discovered motif modules are the binding patterns of TFs and their TF cofactors that frequently bind to the HIF1A ChIP-seq peak regions, which thus represent HIF1A and its cofactor motifs.

**Figure 1.**
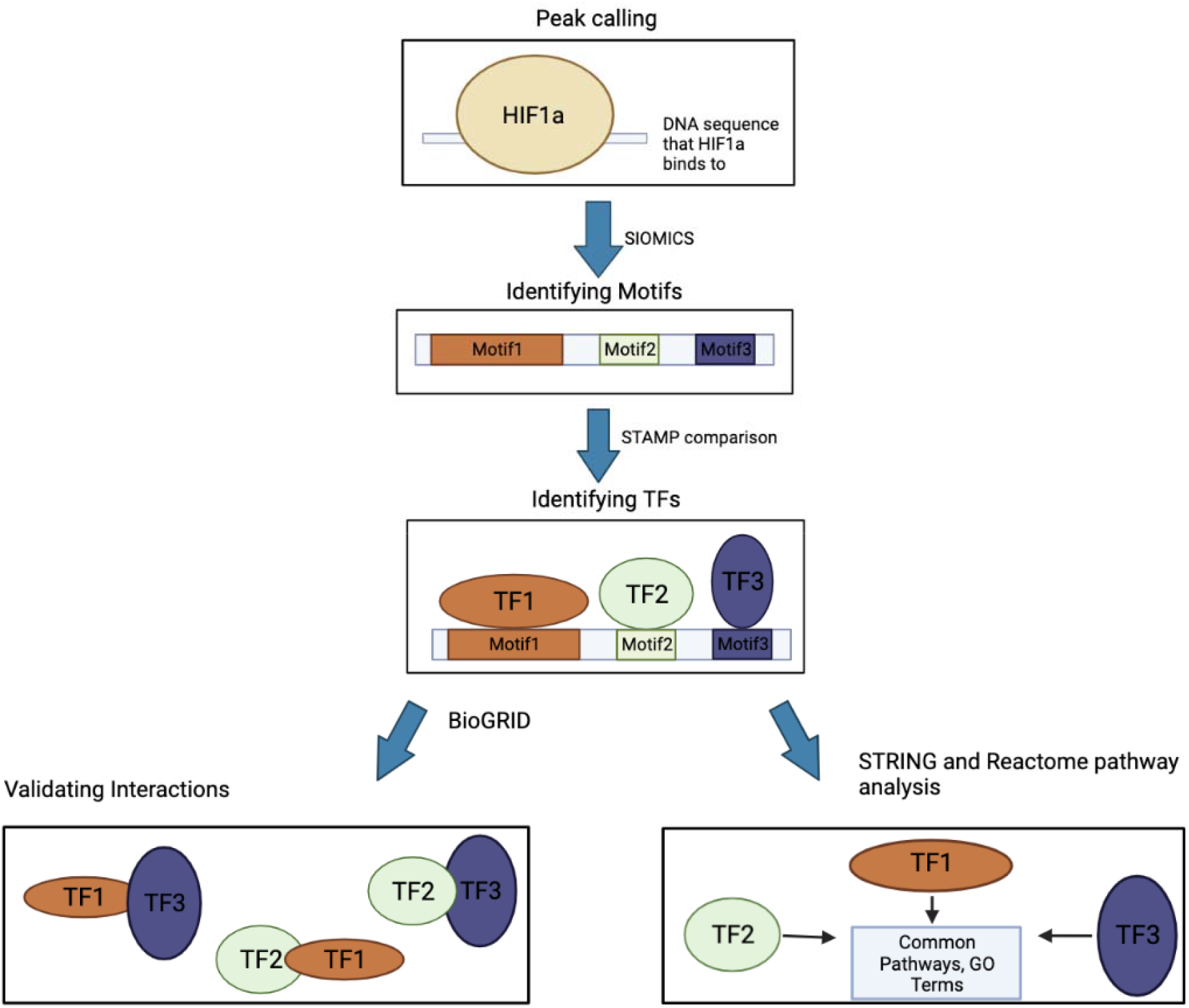
The pipeline to study hypoxia motifs and cofactor pairs.

With the identified motifs in motif modules, we compared the predicted motifs with the known motifs in the JASPAR database ^37^. We claimed that a predicted motif was similar to a known motif if their STAMP comparison E-value was smaller than 1.0E-5 ^38^. A predicted motif similar to a known motif suggests that the TF corresponding to this known motif may play a regulatory role in the data. As a side product, we considered this corresponding TF the HIF1A TF cofactor that binds to this predicted motif. In this way, we obtained 201 HIF1A cofactors.

### Validating the predicted motifs and cofactors

To validate the predicted motifs and cofactors, in addition to comparing the predicted motifs with known motifs, we compared the predicted motifs across the eight cell lines (Supplementary Table S2). Two predicted motifs were similar if they had a STAMP E-value smaller than 1.0E-8. This more stringent E-value cutoff was used as previously ^39^ because these motifs were predicted by the same tool and might be intrinsically more similar. A more stringent cutoff than the comparison of predicted with known motifs may control the false positives better.

Similarly, we compared the predicted motif pairs across cell lines. Recall that SIOMICS outputs motif modules, and a motif module is a group of motifs that co-occur in a significant number of input sequences. We considered all motif pairs in every motif module as the motif pairs in each cell line. Two predicted motif pairs from two cell lines were similar if the corresponding predicted motifs in the two pairs were similar (STAMP E-value < 1.0E-8).

In our study, motif pairs are significant since if the TFs are indeed cofactors of HIF1A, they should be interacting with one another. This combinatorial interaction of TFs within the TF complex is crucial to gene expression. Thus, with the predicted motif pairs, we investigated whether their corresponding cofactor pairs were enriched with known interacting TF pairs in BioGRID V4.4.209 ^11^. BioGRID is a database that stores curated PPIs, including TF-TF interactions. We gathered these TF-TF interactions and split them into two categories – direct and indirect. A direct interaction meant that the two TFs physically interacted, and an indirect interaction indicated that the TFs interacted through a third protein. Following this procedure, we gathered 6904 direct and 101,455 indirect TF-TF interactions. The predicted TF pairs that correspond to the predicted motif pairs in motif modules were then compared with these curated TF-TF interactions in BioGRID in enrichment analysis. Since each motif could correspond to multiple TFs, all of which had STAMP E-value smaller than 1.0E-5, we obtained the corresponding TF pairs from the predicted motif pair in two ways. One was to consider only the TF with its motif most similar to a predicted motif as the TF cofactor behind this predicted motif. The other was to consider up to the top five TFs with their motifs similar to a predicted motif as the TF cofactors of this predicted motif, because the default STAMP outputs only up to the top five TFs. For each of the obtained two sets of predicted cofactor pairs, the enrichment of known TF pairs in the set of predicted cofactor pairs in every cell line was carried out by the hypergeometric testing described in the next section.

Finally, we studied the predicted HIF1A cofactors. We compared the HIF1A direct cofactors with the known HIF1A-interacting-TFs in BioGRID ^11^, HPRD ^12^, and BIND ^13^, obtained from the NCBI page of the HIF1A gene (https://www.ncbi.nlm.nih.gov/gene/3091). Using PubMed, we also searched for additional HIF1A cofactors that were not found in these databases by looking at if a predicted cofactor and HIF1A interact with each other, creating a high likelihood of a regulatory cascade. We used various experiments in the literature to further analyze and verify the predicted HIF1A cofactors.

### Testing the enrichment of known interacting TF pairs in the predicted ones

We tested the enrichment of known interacting TF pairs in the predicted ones by hypergeometric testing. Assume there were *N* TFs and *M* interacting TF pairs in the database. Assume there was *n* of the *N* TFs in a given group of predicted cofactor pairs in a cell line, and *m* cofactor pairs were known to interact. The p-value of the enrichment of the known interacting TF pairs was then calculated as *phyper* 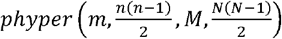 where *phyper* 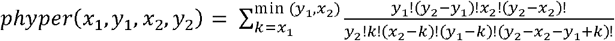 for any non-negative integers x_1_, y_1_, x_2_ and. y_2_^40,41^.

### Gene Ontology (GO) and pathway analysis

To study the regulatory mechanisms of the identified cofactors and the impacts of their cooperativity with HIF1A, we analyzed the binding specificity of the cofactors and their common target genes. We identified the target genes using the annotatePeaks tool from HOMER ^42^ with the GENCODE ^43^ annotation files and assigned every ChIP-seq peak to a gene according to the peak ‘s distance to the nearest transcription start site (TSS). Using the GO and Reactome pathway analysis offered by STRING ^44^, the identified target genes and TFs were then researched regarding their roles in hypoxia-related pathways. Moreover, the tool ChIPseeker ^45^ was also used to analyze the data, which allowed us to visualize the distance of the ChIP-seq peaks to the nearest gene and the pathway enrichment analysis of the ChIP-seq peaks.

## Results

### The predicted motifs and TF cofactors are biologically meaningful

Using the process shown in Figure 1, we could identify motifs and TF cofactors in every cell line (Supplementary Table S1). To see whether the predicted motifs and TF cofactors are biologically meaningful, we compared the predicted motifs with known motifs in the JASPAR database ^37^. In every cell line, on average, approximately 65.6% (Supplementary Table S2) of the predicted motifs were similar to known motifs (STAMP E-value < 1.0E-5) (Table 1). Not every predicted motif has a similar known JASPAR motif because JASPAR does not have known motifs for many TFs, and the current tools to compare motif similarity are imperfect ^36,46^. When we lowered the cutoff to 1.0E-4, more than 83.8% of the identified motifs were similar to a known motif. The high percentages of the predicted motifs similar to the known motifs indicate that these predicted motifs are biologically sound.

**Table 1.** The predicted motifs in eight cell lines.

| Cell lines | #peaks | #motifs | % motifs similar to known motifs (STAMP E-value $\leq 1.0E-5$) | % motifs similar to known motifs (STAMP E-value $\leq 1.0E-4$) | % motifs in other cell lines | % Peaks overlapped across cell lines |
| --- | --- | --- | --- | --- | --- | --- |
| 501A | 13061 | 100 | 60/100 = 60.0% | 81/100 = 81.0% | 90/100 = 90.0% | 66.2% |
| MDA-MB-231 | 1500 | 32 | 25/32 = 78.1% | 29/32 = 90.6% | 32/32 = 100% | 94.1% |
| HepG2 | 12600 | 100 | 68/100 = 68.0% | 88/100 = 88.0% | 97/100 = 97.0% | 60.9% |
| HKC8 | 391 | 100 | 58/100 = 58.0% | 77/100 = 77.0% | 71/100 = 71.0% | 97.2% |
| HUVEC | 661 | 27 | 13/27 = 48.1% | 20/27 = 74.1% | 25/27 = 92.3% | 59.3% |
| K562 | 4156 | 100 | 71/100 = 71.0% | 88/100 = 88.0% | 90/100 = 90.0% | 58.7% |
| LNCaP | 1563 | 98 | 58/98 = 59.2% | 77/98 = 78.6% | 75/98 = 76.5% | 69.1% |
| PC3 | 39025 | 100 | 82/100 = 82.0% | 93/100 = 93.0% | 94/100 = 94.0% | 66.2% |

We also compared the predicted motifs across cell lines. The rationale for this comparison is that if a motif is a bona fide one, it is likely to occur in other cell lines, as different cell lines may share certain regulatory pathways. We found that, on average, approximately 88.9% of the predicted motifs in each cell line were independently identified across cell lines (STAMP E-value < 1.0E-8). This high percentage of motif conservation across cell lines corroborates the biological significance of the predicted motifs and cofactors.

Note that such a high percentage of motif conservation across cell lines was not due to sharing ChIP-seq peaks by cell lines. We observed that the overlap percentage of ChIP-seq peaks did not correlate with the percentage of similar motifs between cell lines. For instance, although more than 87.7 % of peaks in HKC8 overlapped with those in HepG2, only 25.9% of the motifs were similar between the two cell lines. Moreover, we removed the overlapping peaks in two cell lines and still observed a large percentage of shared motifs. For instance, the two cell lines, 501A and HepG2, which had a similar number of ChIP-seq peaks, shared 40.1% of their HIF1A ChIP-seq peaks and 83.0% of the predicted motifs. After removing all overlapping peaks, 59.0% of the motifs in 501A were similar to those in HepG2, and 52.0% of the motifs in HepG2 were similar to those in 501A. The substantial number of motifs still similar after removing overlapping peaks demonstrated that the sharing of the motifs was not due to overlapping peak regions across cell lines. It also suggested that the predicted motifs and cofactors were biologically meaningful.

Finally, we compared the predicted motif pairs across cell lines. More than 63.9% of the motif pairs were independently discovered in other cell lines. To see the significance of such a high percentage of motif pairs independently discovered across cell lines, we generated the same numbers of random motif pairs in each cell line. We randomly chose the predicted motifs in each cell line to form the same number of random motif modules, with each random motif module composed of the same number of motifs as the corresponding predicted motif module. We then similarly obtained random motif pairs from these random motif modules. We observed that a much lower percentage of random motif pairs occurred in more than one cell line (Supplementary Table S2, p-value < 2.1E-15).

### Databases and literature support the predicted cofactor interactions

To evaluate the biological significance of the predicted cofactors, we studied whether the predicted cofactor pairs were enriched with known interacting TF pairs in BioGRID. In brief, for every predicted motif module in each cell line, we obtained the corresponding cofactor pairs in two ways, with every motif pair in each motif module corresponding to one or multiple cofactor pairs (Material and Methods). For the set of all cofactor pairs in each cell line, we then applied hypergeometric testing to see whether the known interacting TF pairs in BioGRID were significantly overrepresented in this set of the predicted cofactor pairs. We found that the known interacting TF pairs were significantly enriched in the predicted ones in all cell lines except HUVEC (Figures 2A, 2B, and Supplementary Table S3). As discussed in the following sections, many interacting TF pairs buried in literature were not curated in BioGRID, such as KLF4, EGR1, NFE2L2, TFAP2A, etc. In this sense, one should consider the above enrichment significance being underestimated. We did not identify any known cofactor pair in HUVEC, likely because HIF1A is not a dominant hypoxia-related TF here, as discussed in the next section.

**Figure 2.**
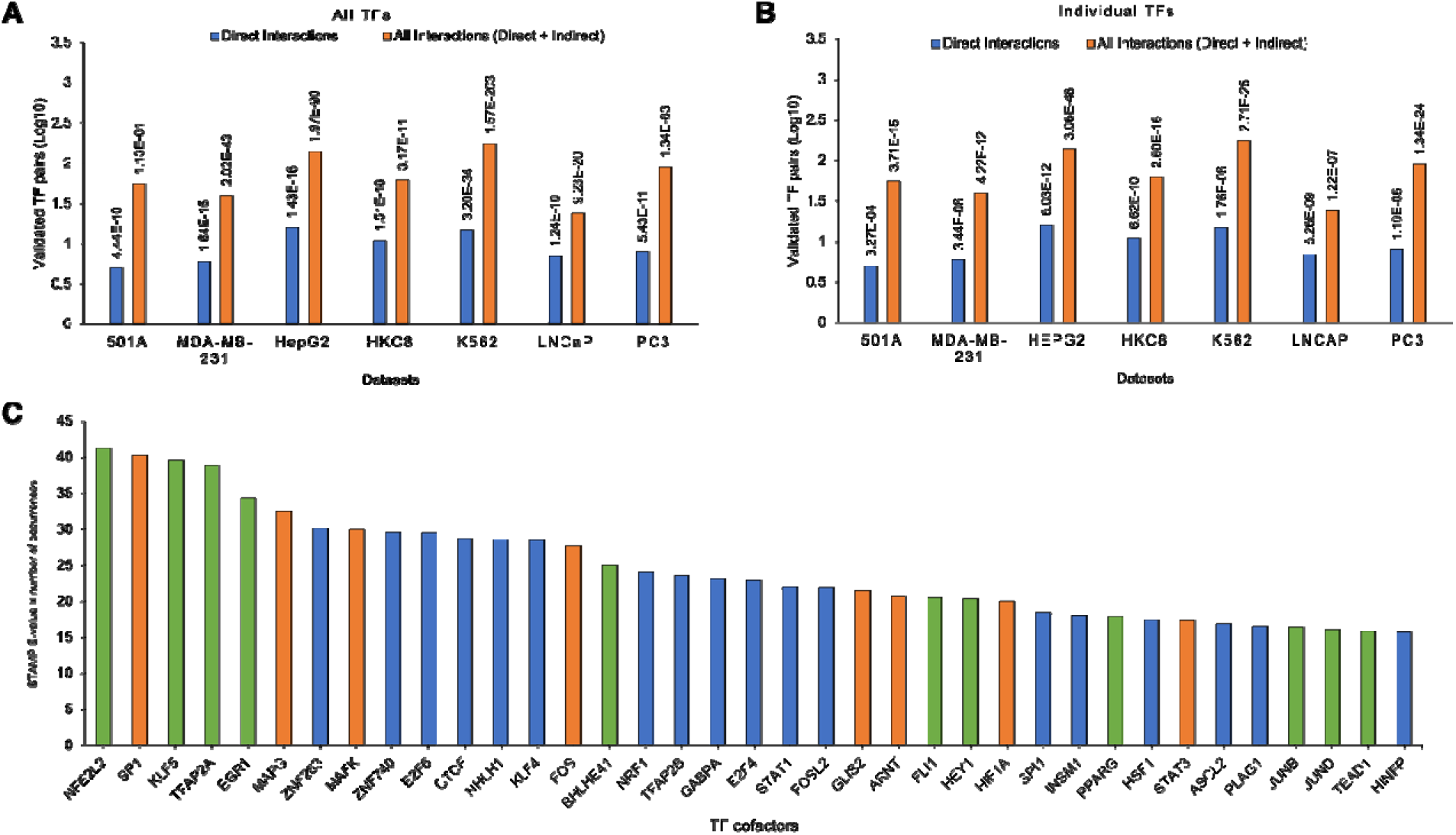
The predicted cofactors are biologically sound. (A) and (B) overrepresentation of the known TF-TF interactions in the predicted cofactor pairs. One predicted motif may correspond to multiple TFs in (A) while only one TF in (B). (C) The top 37 HIF1A cofactors ranked by the multiple of the −log of the STAMP E-value of its predicted motif and the number of motif modules in which this predicted motif occurred. An orange bar indicates that these cofactors were in BIND, HPRD, or BioGRID. A green bar indicates that they were not in the above databases but supported by literature. A blue bar indicates the remaining novel cofactors.

The above analysis was for all predicted cofactor pairs in a cell line. We next sought to study how comprehensively the predicted cofactors included known TFs interacting with HIF1A. We obtained the curated known HIF1A cofactors from BioGRID ^11^, HPRD ^12^, and BIND ^13^, large databases containing many experimentally verified PPIs. We filtered out the curated cofactors that were not TFs using the GO database ^47^ and those that did not have a known motif in JASPAR. In this way, only 49 of the 431 curated cofactors in these databases are TFs, and only 29 of the 49 cofactors have motifs in JASPAR. We found that the predicted cofactors included 21 of these 29 (72.4%) curated cofactors. We also compared the predicted HIF1A cofactors with those in another study by Semenza ^48^ that listed 20 HIF1A TF cofactors, many of which differed from the TFs gathered from BioGRID, HPRD, and BIND. We predicted 15 of the 20 (75.0%) HIF1A cofactors (STAMP E-value <1.0E-5).

We next investigated why eight of the 29 known cofactors were missed. Two cofactors, RORA (STAMP E-value 2.9E-3) and ELK1 (STAMP E-value 4.5E-04), were identified in HKC8 and PC3, respectively, while excluded because they did not meet the STAMP E-value cutoff of 1.0E-5. All of the remaining six cofactors were paralogous to the predicted cofactors. Similarly, two of the twenty cofactors from Semenza did not satisfy the STAMP E-value cutoff, and all of the remaining three cofactors were paralogous to our predicted cofactors.

Although the motifs of paralogous TFs are highly similar, there are subtle differences. The STAMP tool may have recognized such differences and predicted that the known motifs of only certain paralogs are similar to a predicted motif. Furthermore, the default STAMP analysis outputs only up to the top 5 TFs with their known motifs similar to a predicted motif, which may prevent discovering these missed paralogous cofactors. When we allowed STAMP to output more top TFs, all missed paralogous cofactors were discovered. For instance, the six of the 29 known cofactors we missed, CEBPA, TP63, TP73, RUNX2, RUNX3, and ESRRB, had a STAMP E-value of 7.0E-5, 1.7E-4, 1.4E-3, 2.9E-3, and 1.6E-3, respectively. They were missed because their STAMP E-value did not satisfy our cutoff, and they were not in the top five TFs with motifs similar to the corresponding predicted motifs. Interestingly, the missed paralogous TFs might not be involved in the eight cell lines we considered here. We analyzed the experiments where the known interaction was gathered and found that the cell lines used to verify their interaction with HIF1A differed from the ones we used. For instance, to the best of our knowledge, RUNX3 has only been demonstrated in human gastric cells and RUNX3 transfected cells to interact with HIF1A (54), and CREB3L1 has a high tissue-specific expression and is preferentially expressed in only osteoblasts and astrocytes (57), which are thus not validated in the eight cell lines we considered under the hypoxia conditions.

Because the number of the predicted HIF1A-interacting cofactors was large, we focused on the top 37 predicted cofactors that had their motifs co-occur with the HIF1A motif in motif modules (Figure 2C). Intuitively, a top cofactor should have its motif more similar to a known motif and be active in more cell lines. We thus ranked the predicted cofactors based on the product of the −log of the STAMP E-value of its predicted motif and the number of cell lines in which its motif was predicted. A cutoff of 15 was used because a top cofactor should have an average STAMP E-value smaller than 1.0E-5 (Materials and Methods) and occur in more than three of the eight cell lines. We obtained the top 37 cofactors with this cutoff of 15. We found that 19 of these 37 cofactors were directly supported by literature and the PPI databases. This high success rate in discovering known HIF1A cofactors supports the validity of the study and suggests that the predicted cofactors are likely biologically meaningful. Surprisingly, of these 19 cofactors, 12 (63.1%) cofactors were not reported in the current PPI databases (Figure 2C), suggesting that the current PPI databases are far from complete. For instance, four of the top five cofactors – NFE2L2 ^51^, KLF5 ^52^, TFAP2A ^53^, and EGR1 ^54^ - have all been experimentally identified to interact with HIF1A while not being found in the PPI databases. For the remaining 18 novel cofactors, as shown in the last section, we could find experimental evidence to support the direct interaction of HIF1A with at least 11 cofactors. In other words, at least 30 of the top 37 cofactors are validated or supported in the literature.

### HIF1A and cofactors regulate significant cancer-related pathways

After validating the statistical and biological significance of the cofactors, we next sought to identify the mechanisms behind their contributions to hypoxia pathways, especially across different cell lines. We assigned the closest gene to each HIF1A ChIP-seq peak and analyzed the regulation of those genes and their relation to the cofactors (Supplementary Tables S4-S6).

The regulatory regions, especially enhancers in mammalian genomes, are often far from their target genes ^55–58^. We thus first investigated how far the HIF1A ChIP-seq peaks are relative to their closest genes and whether it is proper to assign the closest genes as the target genes of the peak regions. Using the Homer tool ^42^, annotePeaks.pl, we labeled all HIF1A ChIP-seq peaks by analyzing their distances to the nearest TSS. A representation of the binding specificities was then gathered using the tool ChIPseeker (Figure 3). We observed that HIF1A and its respective TF cofactors have binding sites that are mostly found to be clustered around the TSS (Figure 3), showing that HIF1A and most of the TF cofactors identified regulate gene expression by binding to promoters and are primarily promoter-centered TFs, a trend supported by other literature ^19,59^. Most HIF1A ChIP-seq peaks are thus close to the protein-coding genes and likely to regulate the closest protein-coding genes in all cell lines except HUVEC.

**Figure 3.**
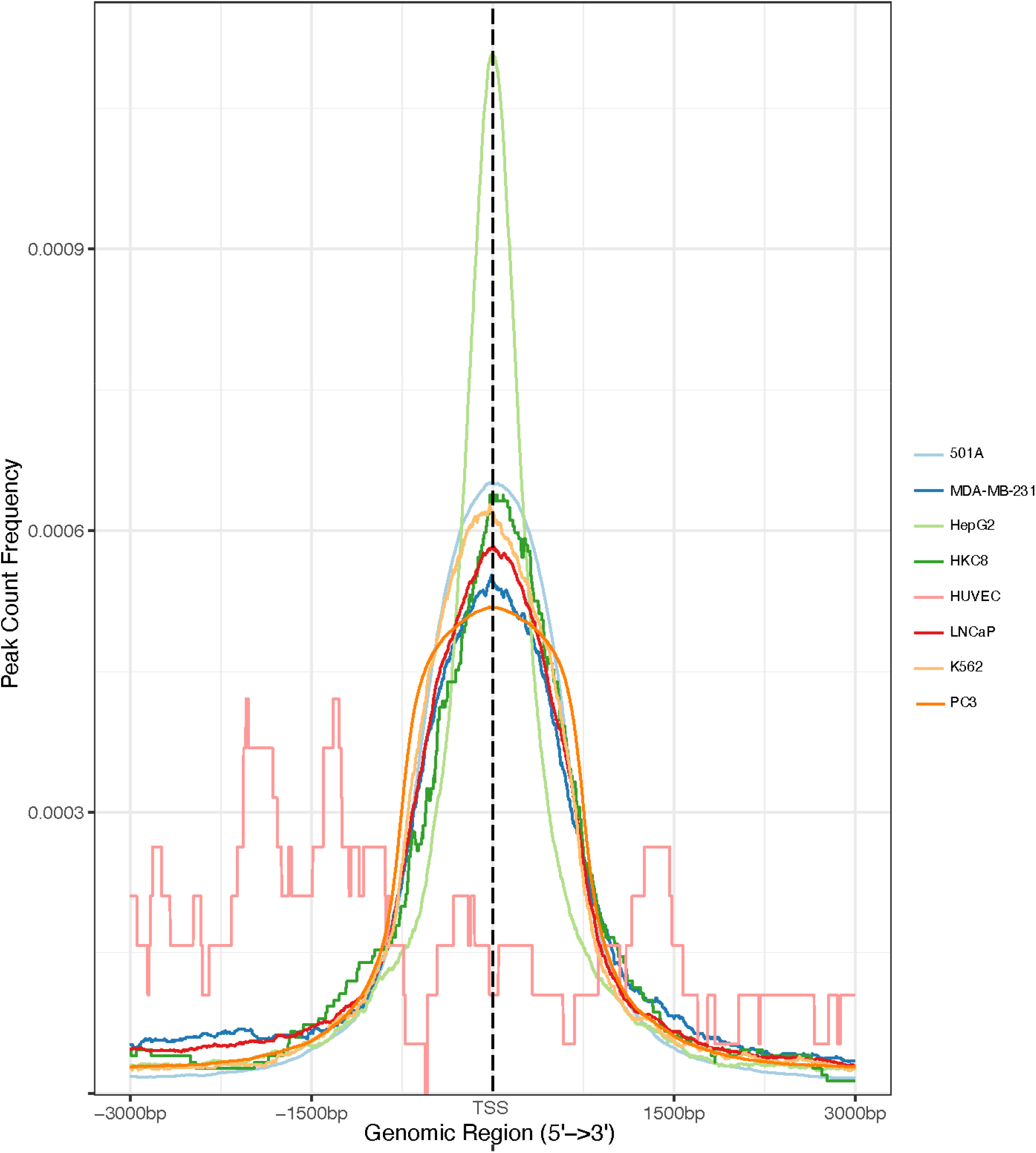
The distance of the HIF1A ChIP-seq peak regions to the proximal genes. Using the ChIPseeker tool, the HIF1A peaks were visualized in terms of their proximity to nearby genes.

With the closest genes as the target genes, we next sought to identify pathways through GO, Reactome, and ChIPseeker pathway analysis (Supplementary Tables S4-S6). We discovered that among the most overrepresented GO terms and pathways, several of them identified were heavily related to hypoxia-induced pathways in cancers. For instance, GO terms like canonical glycolysis, glucose metabolism, IRE1-mediated unfolded protein response ^60^, peptidyl-proline hydroxylation to 4-hydroxy-1-proline ^4^, RHO GTPase cycle ^61^, transcriptional regulation by TP53 ^62^, and cellular response to hypoxia were identified, while similar processes were found in the Reactome and ChIPseeker pathway analysis – transcriptional regulation by TP53, glycolysis, cellular responses to stress, etc. Consistently with all three methods, we found no enriched biological processes in HUVEC.

Surprisingly, HUVEC differs greatly from the rest of our cell lines. As shown in Figure 3, we also noticed that HIF1A’s binding distributions differed from those of other cell lines. One possible explanation is that endothelial PAS domain protein 1 (EPAS1, also known as HIF2A? instead of HIF1A is characterized as the predominant hypoxia inducible factor isoform in HUVEC ^20^. However, it is still uncharacteristic that HIF1A does not have some other regulatory role. Instead, we propose that this is due to HIF1A’s lack of cofactors in the HUVEC cell line. It has been well documented that the cofactors involved in a complex are crucial to binding specificity and gene regulation. Consistent with our claim, according to Table 1, we can see that HUVEC has a low cofactor discovery rate, and in Figure 2, there is a low enrichment for the TF pairs in HUVEC. As a result, we hypothesize that it is due to the lack of cofactors that HIF1A’s binding specificity is different and gene enrichment is minimal. Thus, this further supports the validity of the discovered cofactors in our study and the crucial roles they play in HIF1A’s functions.

We next analyzed how different TF cofactors play different roles in different cell lines, resulting in potential cell-type-specific expression. Despite a high number of cofactors identified across cell lines, the identified cofactors were quite distinct when comparing two cell lines. On average, more than 71.1% of the cofactors in each cell line were not found in another cell line (Supplementary Table S7). Since the difference in cofactors is essential to differential gene expression, we expect that the more different the cofactors are, the more different the enriched GO terms are. To quantify this, we graphed the difference in GO terms against the difference in cofactors in every pair of cell lines (Supplementary Figure S1). We found that for most of the cell lines, a high difference between TF cofactors resulted in a high difference in the enriched GO terms (average R^2^ = 0.77) in all cell lines except MDA-MB-231 (R^2^ = 0.53) and HKC8 (R^2^ = 0.23). This conclusion still held when we compared the difference in the identified pathways with the difference in the cofactors in pairs of cell lines. For instance, through the ChIPseeker pathways, although only 6.7% of the cofactors in MDA-MB-231 and 3.5% of the cofactors in HKC8 were shared (Supplementary Table S7), four of the five most enriched pathways in HKC8 and four of the five most enriched pathways in MDA-MB-231 were identified in both cell lines (Supplementary Table S6). The above analysis suggested that while different TF cofactors help promote various transcriptional responses, HIF1A may also cooperate with different TF cofactors in different cell lines to regulate overlapping target genes and pathways.

### The novel HIF1A cofactors make biological sense

We gathered the leftover TF cofactors that were not supported by external evidence, as found in Figure 2C. In this study, we labeled a predicted cofactor as a probable HIF1A cofactor if it is confirmed to interact with HIF1A directly or indirectly, controlling the expression of HIF1A and vice versa and possess a high similarity in their target genes. We could not go into details about the 201 HIF1A cofactors discovered (Supplementary Table S8), including those in Figure 2C. Instead, we decided to validate the top TF cofactors (Figure 4A) that were not in the databases or directly supported by literature, namely: heat shock transcription factor 1 (HSF1), E2F transcription factor 4 (E2F4), E2F6, nuclear respiratory factor 1 (NRF1), FOS like 2, AP-1 transcription factor subunit (FOSL2), signal transducer and activator of transcription 1 (STAT1) and Kruppel like factor 4 (KLF4) (Figure 4B). For the remaining top cofactors, we filled the supporting evidence as HIF1A cofactors to the best of our knowledge in Supplementary Table S9.

**Figure 4.**
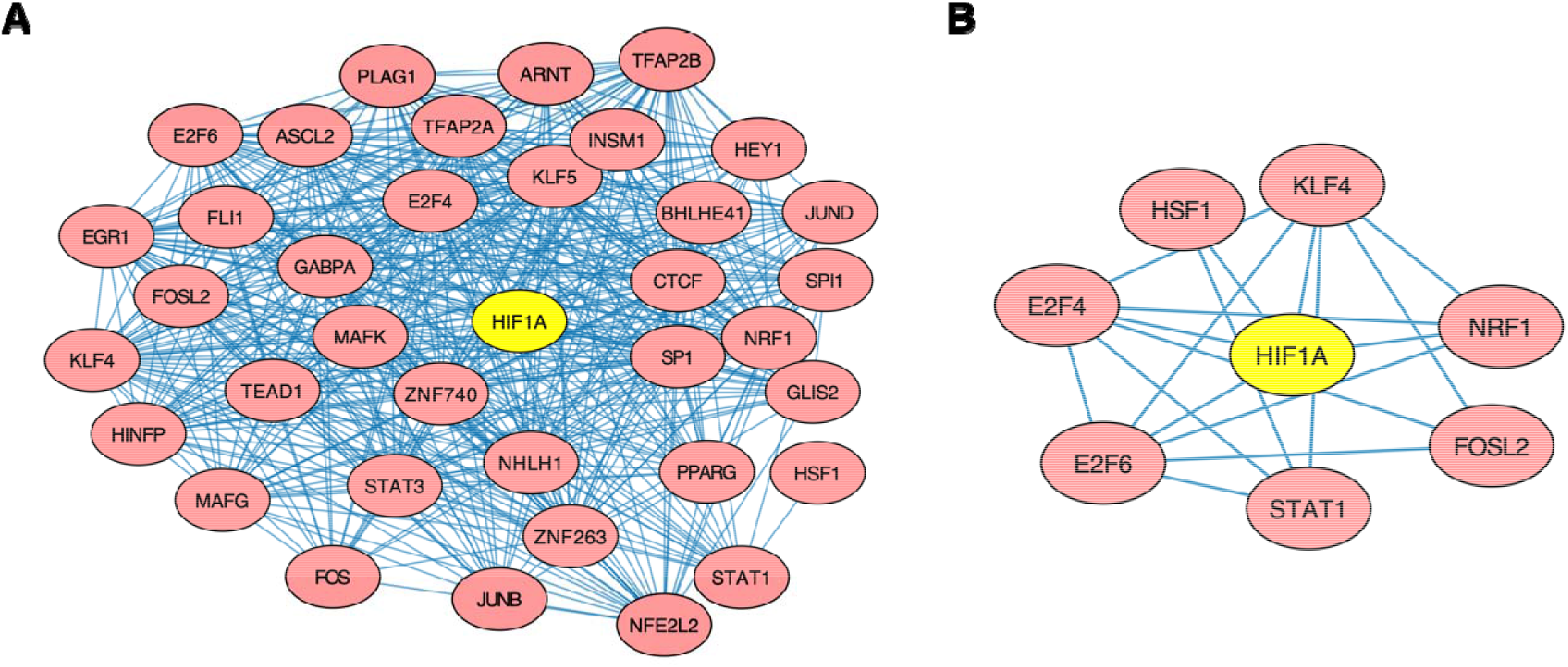
Interactions between discovered HIF1A cofactors. (A) Cytoscape representation of the top 37 TFs and their interactions with one another. (B) Cytoscape network of interactions between six top novel TF cofactors, HSF1, E2F4, E2F6, STAT1, FOSL2, NRF1, and KLF4. SIOMICS predicted the edges shown.

HSF1 is the main human protein responsible for regulating responses to severe heat and inflammation. It activates heat shock proteins like HSP70 and HSP90, which plays an essential role in tumorigenesis, as they allow cancer cells to manage their elevated heat and stress levels resulting from their rapid proliferation ^63^. Due to the connection between hypoxia and cancer, it has also been demonstrated that HSF1 and HIF1A are closely related to tumor formation ^64^. HSF1 regulates the mRNA binding protein, HuR, which controls the expression of HIF1A, and VEGF, a protein that stimulates neovascularization and plays an essential role in responses to hypoxia. A lack of HSF1 has been shown to dramatically deplete HIF1A and decrease angiogenesis, which is directly linked to a cancer cell’s growth and development. Our analysis further supports the importance of HSF1 in hypoxia-related pathways as the GO term “cellular responses to heat stress” was found in all cell lines except HUVEC. Interestingly, though it could mistakenly be inferred that HSF1 is a transcriptional regulator of HIF1A, there are very complex relations between the two TFs. It was found that while HIF1A upregulates HSF1 expression in *Drosophila* cells to activate the heat shock response pathway, HSF2 and HSF4 were found to upregulate HIF1A in mammalian cells ^64^ to activate a hypoxic one. Moreover, we found that HSF1 was ranked as a top ten predicted cofactor by the tool PIP ^65^, and seven of the top nine predictions by which were experimentally verified by previous studies. PIP predicts a potential interacting cofactor by testing the co-expression of the two proteins, orthology, domains, transitive scores, and other factors. All the above evidence shows the interconnectedness of the two TFs, which demonstrates a high possibility of HSF1 engaging in a PPI with HIF1A.

E2F4 and E2F6 are members of the E2F family of TFs and have a high potential to interact with HIF1A directly. They are responsible for regulating cell cycle ^66^. GO terms like “mitotic DNA damage checkpoint, regulation of transcription from RNA polymerase II promoter in response to hypoxia, and cell cycle phase transition” were found in our enrichment analysis (Supplementary Table S5). Extensive studies have also linked both TFs and their functions to hypoxia-related pathways in cancers. For instance, E2F4 was found to work cooperatively with MYC and NRF1 to promote tumorigenesis in hypoxia-driven apoptosis ^67^. Furthermore, Tiana et al. hypothesized that HIF1A might directly interact with E2F4/E2F6 to regulate the activity of SIN3A, a protein essential for a cell’s complete response to hypoxia ^20^. E2F7, a paralog of E2F4/E2F6, has also been found to interact physically with HIF1A to form a repressor complex ^68^.

Interestingly, in the E2F4-NRF1-MYC complex mentioned above, NRF1 was also identified as a potential cofactor, while MYC is a known TF cofactor of HIF1A ^69^. A previous study showed that NRF1 was found to be a regulator of HIF1A in HEK293T cells. Furthermore, Wierenga et al. performed a motif search on HIF1A binding sites ^70^ and found that NRF1 was significantly enriched alongside SP1 ^71^ and ELK1 ^72^, both of which are known HIF1A cofactors. NRF1 has also been discovered to be a pioneer TF ^73^. Pioneer TFs remodel their nearby landscape to allow non-pioneer TFs to interact with a specific DNA sequence, and in this case, HIF1A may interact with NRF1 to increase binding specificity and so that it can regulate target genes that it did not previously have access to.

Much evidence supports the rest of the possible TF cofactors that were identified. For instance, FOSL2 is another potential cofactor. FOSL2 and other members of the FOS family usually dimerize with proteins of the JUN family to form the AP-1 complex ^74^. Our analysis identified many components of the AP-1 complex, including JUN, JUND, JUNB, FOS, FOSL1, and FOSL2. The AP-1 complex has conclusively been shown to interact with HIF1A to express a variety of hypoxia inducible factor target genes, including VEGF, the master regulator of angiogenesis. FOSL2 is also shown to have bound to several of HIF1A’s target genes and is itself upregulated by HIF1A ^75^. Thus, it is very probable that FOSL2 is a possible HIF1A cofactor, either through interacting with HIF1A through the AP-1 complex or as an individual TF. STAT1 is another significant possible HIF1A cofactor identified in this study. While HIF1A has been shown to repress STAT1 ^76^, STAT1 has been, in turn, shown to repress HIF1A ^77^. Furthermore, it has been well documented that STAT1 and STAT3 are closely related in terms of the pathways that they regulate, to the point where interfering with the expression of one protein may indirectly affect the expression of the other ^78^. Since STAT3 is a known TF cofactor of HIF1A ^79^, we predict that since the two TFs are so closely related, both may engage in an interaction with HIF1A. STAT1 and FOSL2 were found in the PIP database due to a high possibility of co-expression, many similar domains, and several common cofactors. Finally, KLF4 also shows high potential as, like NRF1, KLF4 additionally acts as a pioneer TF. It has been shown that while HIF1A upregulates KLF4 in vascular smooth cells ^80^, KLF4 was found to inhibit HIF1A in Huh7 and HepG2 cells ^81^. This high interconnectivity between the two TFs demonstrates a high possibility of them engaging in crosstalk or a regulatory loop.

## Discussion

In this study, we sought to study HIF1A cofactors and how they connect to the role of HIF1A as a master regulator of a cancer cell’s response to hypoxia. Through the analysis of ChIP-seq data in eight cancer cell lines, we identified 201 potential HIF1A cofactors. Most of these identified cofactors were likely to be meaningful, as they were independently identified in other cell lines, enriched with known interacting TF-pairs, supported by curated databases and literature, and were connected to HIF1A’s gene regulatory pathways.

We compared the predicted cofactors with the known HIF1A cofactors curated in public databases. We identify 21 of the 29 known TF cofactors. Two cofactors were missed due to the STAMP E-value cutoff used, suggesting the limitation of the current motif comparison tools. The other six missed cofactors are paralogous TFs to the identified cofactors, likely inactive in the eight cell lines we considered. For instance, the missed cofactor TP63 is paralogous to the identified cofactor TP53. The expression of TP63 was undetectable under hypoxic conditions ^82,83^. The high percentage of the identification of the known HIF1A cofactors suggests that our pipeline to identify cofactors is reliable, and the predicted HIF1A cofactors are biologically meaningful and worth further experimental validation.

We also examined the cofactors not curated in the PPI databases. We found that many of them were supported by the literature. For instance, of the top five cofactors that were not curated in the PPI databases, four of them – NFE2L2, KLF5, TFAP2A, and EGR1 – had been experimentally verified as HIF1A cofactors in the literature. We also discovered that more than 51.3% of the top 37 cofactors were directly supported by experimental evidence in literature. We analyzed the remaining top cofactors that were not curated in databases or had their interaction with HIF1A experimentally validated. We found evidence supporting their interaction with HIF1A to modulate gene expression during hypoxia.

Recent advancements in understanding the PPI’s role in diseases have made targeting PPIs a promising venture, especially in cancer therapies ^84^. Due to the essential role that many of the discovered cofactors play in regulating HIF1A target genes, these new cofactors may have regulatory mechanisms that could be exploited for novel therapies. For instance, TFAP2A, an identified cofactor not found in any of the major databases, was demonstrated to interact with HIF1A to target the VEGF pathway in nasopharyngeal carcinoma cells. Shi et al. showed that inhibiting TFAP2A led to decreased tumor growth and conclusively proved that TFAP2A had the potential to be investigated as a therapeutic target ^53^. Many of the other new potential cofactors identified also have similar implications. For instance, Toth et al. predicted that simultaneously targeting the HIF1A and NFE2L2 presents a novel approach for cancer therapies and proposed several molecular inhibitors that would target both proteins ^85^.

In five of the eight cell lines, we predicted 100 motifs, which was the upper limit when running the default SIOMICS. The discovered 100 motifs suggested that there could be more motifs unreported. In the future, we could explore the remaining motifs and study other cell lines and types. Moreover, to gain a more comprehensive view of the HIF1A cofactors in hypoxia in cancer cells, experimental research like co-immunoprecipitation arrays would help verify our predictions. We look forward to further investigating the effects of HIF1A’s cofactors and the importance of TF cooperativity in many of its pathways.

## Supporting information

Supplementary result

## Acknowledgment

This work was supported by the National Science Foundation (1661414, 2015838, 2120907). Figure 1 was created with the tools at BioRender.com.

## Author Contributions

H.H. and X.L. conceived the idea. Y.Z. implemented the idea and generated results. Y.Z., S.W., H.H., and X. L. analyzed the data and produced the results. Y.Z., H.H., and X. L. wrote the manuscript. All authors reviewed the manuscript.

## Data availability

The human genome used is from https://www.ncbi.nlm.nih.gov/assembly/GCF000001405.26/. The ChIP-seq data in the eight cell lines are from the following publications ^17–23^. The links to these ChIP-seq data are provided in Supplementary Table S1.

