## Supplementary result for "A systematic study of HIF1A cofactors in hypoxic cancer cells"

**Supplementary Figures**

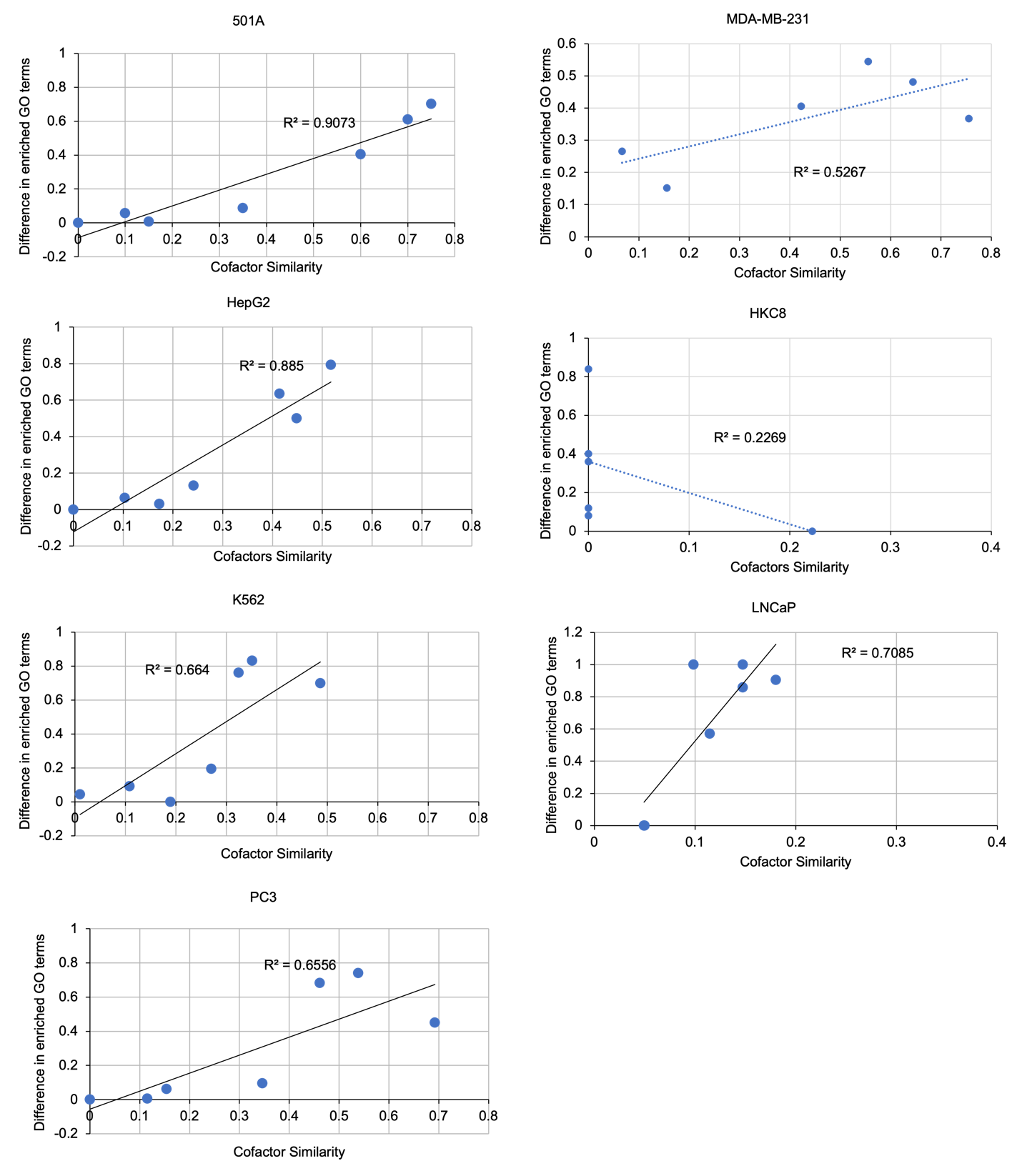
 **Supplementary Figure S1. Correlation between the similarity of cofactors and the similarity in GO terms/genomic processes.** In these graphs, we determined the correlation between the difference in cofactors and the GO terms associated with the cofactors in each cell line. While a linear trend is shown in the 501A, HepG2, K562, LNCaP, and PC3 cell lines, this is not true for the MDA-MB-231 and HKC8 cell lines.

**Supplementary Table S1.** **Sources of the ChIP-seq data as gathered from 8 cell lines**.

| **Dataset** | **Number of peaks identified** | **SRA** | **Links** | **Mean** | **Medium** | **Minimum** | **Maximum** |
| --- | --- | --- | --- | --- | --- | --- | --- |
| 501A | 13061 | SRR5280434 | <https://www.ncbi.nlm.nih.gov/sra/?term=SRX2584126>, | 1172 | 1119 | 800 | 5781 |
| MDA-MB-231 | 1500 | SRR6493909, SRR6493910 | <https://www.ncbi.nlm.nih.gov/sra/?term=SRX3583255>, <https://www.ncbi.nlm.nih.gov/sra/?term=SRX3583256> | 1177 | 1124 | 800 | 5781 |
| HepG2 | 12600 | SRR7968966, SRR7968967 | <https://www.ncbi.nlm.nih.gov/sra/?term=SRX4802361>, <https://www.ncbi.nlm.nih.gov/sra/?term=SRX4802362> | 1077 | 1062 | 227 | 5781 |
| HKC8 | 391 | SRR7968891, SRR7968892 | <https://www.ncbi.nlm.nih.gov/sra/?term=SRX4802306>, <https://www.ncbi.nlm.nih.gov/sra/?term=SRX4802307> | 1016 | 1014 | 227 | 5781 |
| HUVEC | 661 | SRR5018853 | <https://www.ncbi.nlm.nih.gov/sra/?term=SRX2346891>, | 1181 | 1135 | 750 | 5781 |
| K562 | 4156 | SRR10820724, SRR10820725 | <https://www.ncbi.nlm.nih.gov/sra/?term=SRX7494163>, <https://www.ncbi.nlm.nih.gov/sra/?term=SRX7494164> | 906 | 906 | 227 | 5781 |
| LNCaP | 1563 | SRR7192349 | <https://www.ncbi.nlm.nih.gov/sra/?term=SRX4108920> | 902 | 900 | 227 | 5781 |
| PC3 | 39025 | SRR6233822, SRR6233823, SRR6233824, SRR6233825, SRR6233826, SRR6233827, SRR6233828, SRR6233829 | <https://www.ncbi.nlm.nih.gov/sra/?term=SRX3342251>, <https://www.ncbi.nlm.nih.gov/sra/?term=SRX3342252> | 977 | 968 | 227 | 5781 |

There was a minimum of 391 peaks and a maximum of 39025 peaks. The average peak number was 9119.6 peaks, and the median was 2859.5 peaks. The last four columns are the statistics of the length of the MACS2 peak regions. Certain peak regions are shorter than 800 base pairs because they are close to the chromosome boundaries.

**Supplementary Table S2. Significance of the gathered motif pairs.**

| Cell lines | Number of Predicted Motif Pairs | Number of motif pairs identified in other cell lines | percentage of random motif pairs found in other cell lines | p-value of random motif pairs |
| --- | --- | --- | --- | --- |
| 501A | 1762 | 1428/1762= 81.0% | 14.3% | 0 |
| MDA-MB-231 | 221 | 212/221= 95.9% | 35.8% | 0 |
| HepG2 | 2158 | 1730/2158 = 80.2% | 10.7% | 0 |
| HKC8 | 499 | 168/499 = 33.7% | 7.3% | 0 |
| HUVEC | 42 | 14/42= 33.3% | 1.6% | 2.1E-15 |
| K562 | 2207 | 1207/2207= 54.7% | 25.0% | 0 |
| LNCaP | 332 | 181/332 = 54.5% | 18.4% | 2.2E-16 |
| PC3 | 1793 | 1411/1793 = 78.7% | 17.8% | 0 |

Motif pairs were generated from SIOMICS. A motif pair is considered similar to another motif pair if the two motifs in one motif pair are similar (STAMP E-value < 1E-8) to the two motifs in the other pair. The significance is calculated using $1-pbinom\left( k-1, n, p \right),$where k is the number of shared pairs between a cell line and other cell lines, n is the total number of motif pairs predicted in this cell line, and p is the percentage of random motif pairs in this cell line that are also found in other cell lines.

**Supplementary Table S3. Enrichment analysis of the predicted TF pairs using hypergeometric testing**.

| Cell lines | Considering all possible TF | | Considering most matching TF only | |
| --- | --- | --- | --- | --- |
|  | direct interactions | Merged (direct and indirect) | direct interactions | Merged (direct and indirect) |
| 501A | 4.44E-10 | 1.13E-61 | 3.27E-04 | 3.71E-15 |
| MDA-MB-231 | 1.64E-15 | 2.02E-43 | 3.44E-06 | 4.22E-12 |
| HepG2 | 1.43E-16 | 1.97E-90 | 6.03E-12 | 3.06E-48 |
| HKC8 | 1.51E-18 | 3.17E-11 | 6.62E-10 | 2.59E-16 |
| K562 | 3.20E-34 | 1.57E-203 | 1.76E-06 | 2.71E-25 |
| LNCaP | 1.24E-10 | 9.23E-20 | 6.26E-09 | 1.22E-07 |
| PC3 | 5.43E-11 | 1.34E-83 | 1.19E-05 | 1.34E-24 |

Since each motif could correspond to multiple TFs, we performed two tests. One was to consider up to top 5 TFs that corresponded to the motif (STAMP E-value<1.0E-5), and the other was to consider the TF with the most similar motif (STAMP E-value<1.0E-5). We also performed this enrichment analysis using direct and indirect interactions. A direct interaction meant that two TFs were found to physically interact in an experimental setting. An indirect interaction meant that the two TFs would interact through a common third protein. We gathered the direct and indirect TF interactions from BioGRID. There were 1349 TFs that were involved 6904 direct TF pairs, and 1070 TFs that were involved in 101,455 indirect TF pairs. As shown, known interacting TF pairs was significantly enriched in the predicted ones in every cell line.

**Supplementary Table S4.** **Top 25** **Gene Ontology (GO) terms associated with the target genes of each cell line.**

| Cell lines | Term ID | Term description | Strength | False discovery rate |
| --- | --- | --- | --- | --- |
| 501A | GO:0036498 | IRE1-mediated unfolded protein response | 0.3 | 3.38E-02 |
|  | GO:0046148 | Pigment biosynthetic process | 0.3 | 3.94E-02 |
|  | GO:0016241 | Regulation of macroautophagy | 0.26 | 6.53E-05 |
|  | GO:0016482 | Cytosolic transport | 0.26 | 4.50E-04 |
|  | GO:0044773 | Mitotic DNA damage checkpoint | 0.26 | 7.90E-03 |
|  | GO:0042147 | Retrograde transport, endosome to golgi | 0.26 | 1.97E-02 |
|  | GO:0006446 | Regulation of translational initiation | 0.26 | 3.43E-02 |
|  | GO:0044774 | Mitotic DNA integrity checkpoint | 0.25 | 7.70E-03 |
|  | GO:1903747 | Regulation of establishment of protein localization to mitochondrion | 0.25 | 4.03E-02 |
|  | GO:0000077 | DNA damage checkpoint | 0.24 | 1.90E-03 |
|  | GO:0010822 | Positive regulation of mitochondrion organization | 0.24 | 7.30E-03 |
|  | GO:0032436 | Positive regulation of proteasomal ubiquitin-dependent protein catabolic process | 0.24 | 1.85E-02 |
|  | GO:0032006 | Regulation of TOR signaling | 0.24 | 2.58E-02 |
|  | GO:0010506 | Regulation of autophagy | 0.23 | 3.39E-07 |
|  | GO:0017148 | Negative regulation of translation | 0.23 | 5.70E-03 |
|  | GO:2000060 | Positive regulation of ubiquitin-dependent protein catabolic process | 0.23 | 1.45E-02 |
|  | GO:0000070 | Mitotic sister chromatid segregation | 0.23 | 2.42E-02 |
|  | GO:0031570 | DNA integrity checkpoint | 0.22 | 4.30E-03 |
|  | GO:0007093 | Mitotic cell cycle checkpoint | 0.22 | 4.90E-03 |
|  | GO:2000134 | Negative regulation of g1/s transition of mitotic cell cycle | 0.22 | 3.85E-02 |
|  | GO:0018205 | Peptidyl-lysine modification | 0.21 | 4.18E-05 |
|  | GO:0016197 | Endosomal transport | 0.21 | 3.50E-04 |
|  | GO:0032388 | Positive regulation of intracellular transport | 0.21 | 6.00E-04 |
|  | GO:0032869 | Cellular response to insulin stimulus | 0.21 | 5.00E-03 |
|  | GO:0140014 | Mitotic nuclear division | 0.21 | 8.60E-03 |
| MDA-MB-231 | GO:0018401 | Peptidyl-proline hydroxylation to 4-hydroxy-l-proline | 1.23 | 2.90E-03 |
|  | GO:0019511 | Peptidyl-proline hydroxylation | 1.14 | 1.60E-03 |
|  | GO:0006002 | Fructose 6-phosphate metabolic process | 1.12 | 2.38E-02 |
|  | GO:0018126 | Protein hydroxylation | 1.06 | 8.20E-05 |
|  | GO:0061621 | Canonical glycolysis | 1.06 | 8.20E-05 |
|  | GO:0006007 | Glucose catabolic process | 1.05 | 3.97E-05 |
|  | GO:0046031 | ADP metabolic process | 0.98 | 1.37E-06 |
|  | GO:0006096 | Glycolytic process | 0.97 | 3.97E-05 |
|  | GO:0009179 | Purine ribonucleoside diphosphate metabolic process | 0.94 | 6.02E-07 |
|  | GO:0019320 | Hexose catabolic process | 0.93 | 1.17E-05 |
|  | GO:0019674 | NAD metabolic process | 0.92 | 8.20E-05 |
|  | GO:0046939 | Nucleotide phosphorylation | 0.86 | 4.16E-05 |
|  | GO:0006165 | Nucleoside diphosphate phosphorylation | 0.85 | 1.10E-04 |
|  | GO:0006090 | Pyruvate metabolic process | 0.83 | 3.97E-05 |
|  | GO:0019319 | Hexose biosynthetic process | 0.71 | 4.88E-02 |
|  | GO:0018208 | Peptidyl-proline modification | 0.67 | 4.08E-02 |
|  | GO:0061418 | Regulation of transcription from RNA polymerase II promoter in response to hypoxia | 0.65 | 1.88E-02 |
|  | GO:0016052 | Carbohydrate catabolic process | 0.64 | 1.20E-03 |
|  | GO:0043620 | Regulation of DNA-templated transcription in response to stress | 0.63 | 7.70E-04 |
|  | GO:0043618 | Regulation of transcription from RNA polymerase II promoter in response to stress | 0.63 | 1.50E-03 |
|  | GO:0071456 | Cellular response to hypoxia | 0.62 | 1.49E-05 |
|  | GO:0036294 | Cellular response to decreased oxygen levels | 0.61 | 1.17E-05 |
|  | GO:0006006 | Glucose metabolic process | 0.61 | 4.20E-03 |
|  | GO:0019318 | Hexose metabolic process | 0.58 | 9.70E-04 |
|  | GO:0034101 | Erythrocyte homeostasis | 0.58 | 4.29E-02 |
| HepG2 | GO:0061014 | Positive regulation of mRNA catabolic process | 0.36 | 1.46E-02 |
|  | GO:0090503 | RNA phosphodiester bond hydrolysis, exonucleolytic | 0.36 | 3.08E-02 |
|  | GO:0035195 | Gene silencing by miRNA | 0.35 | 3.48E-02 |
|  | GO:0016441 | Posttranscriptional gene silencing | 0.33 | 2.09E-02 |
|  | GO:0035194 | Post-transcriptional gene silencing by RNA | 0.33 | 2.30E-02 |
|  | GO:0032922 | Circadian regulation of gene expression | 0.31 | 2.52E-02 |
|  | GO:0031047 | Gene silencing by RNA | 0.29 | 1.27E-02 |
|  | GO:0017148 | Negative regulation of translation | 0.28 | 1.10E-03 |
|  | GO:1903313 | Positive regulation of mRNA metabolic process | 0.28 | 2.79E-02 |
|  | GO:0034249 | Negative regulation of cellular amide metabolic process | 0.27 | 1.00E-03 |
|  | GO:0061013 | Regulation of mRNA catabolic process | 0.24 | 4.70E-04 |
|  | GO:0043488 | Regulation of mRNA stability | 0.24 | 2.00E-03 |
|  | GO:0043487 | Regulation of RNA stability | 0.23 | 2.20E-03 |
|  | GO:0032434 | Regulation of proteasomal ubiquitin-dependent protein catabolic process | 0.23 | 1.21E-02 |
|  | GO:0007030 | Golgi organization | 0.23 | 2.64E-02 |
|  | GO:0006006 | Glucose metabolic process | 0.23 | 4.15E-02 |
|  | GO:0018205 | Peptidyl-lysine modification | 0.22 | 1.50E-04 |
|  | GO:0022618 | Ribonucleoprotein complex assembly | 0.22 | 5.60E-03 |
|  | GO:0006611 | Protein export from nucleus | 0.22 | 2.29E-02 |
|  | GO:0007623 | Circadian rhythm | 0.22 | 2.37E-02 |
|  | GO:0006839 | Mitochondrial transport | 0.21 | 3.00E-03 |
|  | GO:0032388 | Positive regulation of intracellular transport | 0.21 | 4.30E-03 |
|  | GO:0033500 | Carbohydrate homeostasis | 0.21 | 8.00E-03 |
|  | GO:0071826 | Ribonucleoprotein complex subunit organization | 0.21 | 9.20E-03 |
|  | GO:0042593 | Glucose homeostasis | 0.21 | 9.90E-03 |
| HKC8 | GO:0018401 | Peptidyl-proline hydroxylation to 4-hydroxy-l-proline | 1.73 | 1.10E-03 |
|  | GO:0061621 | Canonical glycolysis | 1.58 | 3.08E-06 |
|  | GO:0006096 | Glycolytic process | 1.47 | 1.69E-06 |
|  | GO:0018126 | Protein hydroxylation | 1.44 | 9.70E-04 |
|  | GO:0046031 | ADP metabolic process | 1.43 | 5.85E-07 |
|  | GO:0019674 | NAD metabolic process | 1.42 | 3.08E-06 |
|  | GO:0009179 | Purine ribonucleoside diphosphate metabolic process | 1.38 | 5.85E-07 |
|  | GO:0006165 | Nucleoside diphosphate phosphorylation | 1.36 | 1.46E-06 |
|  | GO:0006090 | Pyruvate metabolic process | 1.28 | 3.08E-06 |
|  | GO:0006094 | Gluconeogenesis | 1.2 | 7.80E-03 |
|  | GO:0006006 | Glucose metabolic process | 1.01 | 9.00E-04 |
|  | GO:0016051 | Carbohydrate biosynthetic process | 0.97 | 1.40E-03 |
|  | GO:0036294 | Cellular response to decreased oxygen levels | 0.95 | 1.20E-05 |
|  | GO:0071456 | Cellular response to hypoxia | 0.93 | 6.16E-05 |
|  | GO:0046034 | ATP metabolic process | 0.86 | 9.00E-04 |
|  | GO:0005996 | Monosaccharide metabolic process | 0.83 | 4.10E-03 |
|  | GO:0036293 | Response to decreased oxygen levels | 0.82 | 3.20E-06 |
|  | GO:0001666 | Response to hypoxia | 0.81 | 1.20E-05 |
|  | GO:0009150 | Purine ribonucleotide metabolic process | 0.69 | 1.65E-02 |
|  | GO:0006091 | Generation of precursor metabolites and energy | 0.67 | 2.00E-03 |
|  | GO:0005975 | Carbohydrate metabolic process | 0.61 | 7.70E-03 |
|  | GO:0009117 | Nucleotide metabolic process | 0.58 | 4.56E-02 |
|  | GO:0055114 | Oxidation-reduction process | 0.5 | 2.60E-03 |
|  | GO:0009628 | Response to abiotic stimulus | 0.41 | 3.06E-02 |
|  | GO:0006464 | Cellular protein modification process | 0.26 | 3.37E-02 |
| K562 | GO:0061621 | Canonical glycolysis | 0.7 | 6.30E-03 |
|  | GO:1900034 | Regulation of cellular response to heat | 0.6 | 1.32E-05 |
|  | GO:0006734 | NADH metabolic process | 0.59 | 1.40E-02 |
|  | GO:0009179 | Purine ribonucleoside diphosphate metabolic process | 0.56 | 1.40E-03 |
|  | GO:0046031 | ADP metabolic process | 0.56 | 6.80E-03 |
|  | GO:0006096 | Glycolytic process | 0.56 | 2.30E-02 |
|  | GO:0051031 | tRNA transport | 0.56 | 3.50E-02 |
|  | GO:0097064 | ncRNA export from nucleus | 0.55 | 4.00E-02 |
|  | GO:0071479 | Cellular response to ionizing radiation | 0.52 | 3.60E-03 |
|  | GO:0016925 | Protein sumoylation | 0.51 | 6.50E-03 |
|  | GO:0019674 | NAD metabolic process | 0.51 | 4.56E-02 |
|  | GO:2000573 | Positive regulation of DNA biosynthetic process | 0.46 | 2.19E-02 |
|  | GO:0046794 | Transport of virus | 0.46 | 4.78E-02 |
|  | GO:0070301 | Cellular response to hydrogen peroxide | 0.43 | 4.69E-02 |
|  | GO:0006406 | mRNA export from nucleus | 0.39 | 1.82E-02 |
|  | GO:0048024 | Regulation of mRNA splicing, via spliceosome | 0.39 | 2.01E-02 |
|  | GO:0043484 | Regulation of RNA splicing | 0.38 | 3.40E-03 |
|  | GO:0071426 | Ribonucleoprotein complex export from nucleus | 0.37 | 2.15E-02 |
|  | GO:0006405 | RNA export from nucleus | 0.36 | 1.91E-02 |
|  | GO:2000278 | Regulation of DNA biosynthetic process | 0.35 | 4.89E-02 |
|  | GO:0051028 | mRNA transport | 0.34 | 2.27E-02 |
|  | GO:0050684 | Regulation of mRNA processing | 0.34 | 2.42E-02 |
|  | GO:0046034 | ATP metabolic process | 0.33 | 6.20E-03 |
|  | GO:0006611 | Protein export from nucleus | 0.33 | 2.57E-02 |
|  | GO:0051054 | Positive regulation of DNA metabolic process | 0.31 | 1.18E-02 |
| LNCaP | GO:0018205 | Peptidyl-lysine modification | 0.39 | 1.61E-02 |
|  | GO:0044283 | Small molecule biosynthetic process | 0.28 | 4.29E-02 |
|  | GO:1901566 | Organonitrogen compound biosynthetic process | 0.22 | 3.30E-03 |
|  | GO:0044271 | Cellular nitrogen compound biosynthetic process | 0.21 | 1.60E-03 |
|  | GO:1901576 | Organic substance biosynthetic process | 0.18 | 2.50E-04 |
|  | GO:0044249 | Cellular biosynthetic process | 0.18 | 2.80E-04 |
|  | GO:0034645 | Cellular macromolecule biosynthetic process | 0.18 | 2.30E-02 |
|  | GO:0016070 | RNA metabolic process | 0.18 | 4.33E-02 |
|  | GO:0009058 | Biosynthetic process | 0.17 | 2.70E-04 |
|  | GO:0034641 | Cellular nitrogen compound metabolic process | 0.15 | 1.40E-03 |
|  | GO:1901360 | Organic cyclic compound metabolic process | 0.15 | 2.00E-03 |
|  | GO:0006725 | Cellular aromatic compound metabolic process | 0.15 | 4.90E-03 |
|  | GO:0046483 | Heterocycle metabolic process | 0.15 | 6.90E-03 |
|  | GO:0006139 | Nucleobase-containing compound metabolic process | 0.15 | 7.60E-03 |
|  | GO:0044237 | Cellular metabolic process | 0.11 | 4.66E-07 |
|  | GO:0044260 | Cellular macromolecule metabolic process | 0.11 | 2.10E-03 |
|  | GO:0008152 | Metabolic process | 0.1 | 2.07E-05 |
|  | GO:0044238 | Primary metabolic process | 0.1 | 2.20E-04 |
|  | GO:0006807 | Nitrogen compound metabolic process | 0.1 | 4.20E-04 |
|  | GO:0071704 | Organic substance metabolic process | 0.09 | 2.50E-04 |
|  | GO:0043170 | Macromolecule metabolic process | 0.09 | 2.07E-02 |
| PC3 | GO:0007409 | Axonogenesis | 0.12 | 1.08E-02 |
|  | GO:0000209 | Protein polyubiquitination | 0.12 | 2.12E-02 |
|  | GO:0010506 | Regulation of autophagy | 0.12 | 2.44E-02 |
|  | GO:0051056 | Regulation of small GTPase mediated signal transduction | 0.12 | 2.64E-02 |
|  | GO:0097485 | Neuron projection guidance | 0.12 | 3.24E-02 |
|  | GO:0007411 | Axon guidance | 0.12 | 3.37E-02 |
|  | GO:0044770 | Cell cycle phase transition | 0.12 | 4.62E-02 |
|  | GO:0000278 | Mitotic cell cycle | 0.11 | 2.90E-04 |
|  | GO:0031344 | Regulation of cell projection organization | 0.11 | 3.50E-04 |
|  | GO:0120035 | Regulation of plasma membrane bounded cell projection organization | 0.11 | 3.80E-04 |
|  | GO:0000902 | Cell morphogenesis | 0.11 | 4.10E-04 |
|  | GO:1903047 | Mitotic cell cycle process | 0.11 | 4.60E-04 |
|  | GO:0007346 | Regulation of mitotic cell cycle | 0.11 | 8.60E-04 |
|  | GO:0031175 | Neuron projection development | 0.11 | 9.50E-04 |
|  | GO:0051640 | Organelle localization | 0.11 | 1.20E-03 |
|  | GO:0000904 | Cell morphogenesis involved in differentiation | 0.11 | 1.70E-03 |
|  | GO:0032990 | Cell part morphogenesis | 0.11 | 2.20E-03 |
|  | GO:0010975 | Regulation of neuron projection development | 0.11 | 2.60E-03 |
|  | GO:0048858 | Cell projection morphogenesis | 0.11 | 2.60E-03 |
|  | GO:0120039 | Plasma membrane bounded cell projection morphogenesis | 0.11 | 2.70E-03 |
|  | GO:0048812 | Neuron projection morphogenesis | 0.11 | 3.20E-03 |
|  | GO:1902532 | Negative regulation of intracellular signal transduction | 0.11 | 5.80E-03 |
|  | GO:0048667 | Cell morphogenesis involved in neuron differentiation | 0.11 | 6.70E-03 |
|  | GO:0061564 | Axon development | 0.11 | 8.70E-03 |
|  | GO:1901990 | Regulation of mitotic cell cycle phase transition | 0.11 | 1.22E-02 |

This was found by inputting the gathered genes into STRING (2) and performing a GO Enrichment. The enriched GO terms were then sorted by their strength. As observed, several of the top GO terms in each cell line were related to HIF1A mediated pathways. For example, the most enriched GO terms in the MDA-MB-231 cell line were peptidyl-proline hydroxylation to 4-hydroxy-l-proline, which is a process that is significantly regulated by HIF1A (3).

**Supplementary Table S5. Top enriched Reactome pathways associated with the target genes of each cell line.**

| Cell line | Term ID | Enriched Reactome pathways | Strength | False discovery rate |
| --- | --- | --- | --- | --- |
| 501A | HSA-165159 | MTOR signalling | 0.34 | 4.99E-02 |
|  | HSA-68875 | Mitotic Prophase | 0.24 | 2.29E-02 |
|  | HSA-1632852 | Macroautophagy | 0.23 | 1.16E-02 |
|  | HSA-9612973 | Autophagy | 0.21 | 1.54E-02 |
|  | HSA-6811442 | Intra-Golgi and retrograde Golgi-to-ER traffic | 0.2 | 9.60E-03 |
|  | HSA-2559583 | Cellular Senescence | 0.2 | 2.27E-02 |
|  | HSA-3700989 | Transcriptional Regulation by TP53 | 0.19 | 2.40E-04 |
|  | HSA-983168 | Antigen processing: Ubiquitination & Proteasome degradation | 0.19 | 9.80E-04 |
|  | HSA-3247509 | Chromatin modifying enzymes | 0.19 | 3.40E-03 |
|  | HSA-5633007 | Regulation of TP53 Activity | 0.19 | 3.20E-02 |
|  | HSA-1640170 | Cell Cycle | 0.18 | 4.73E-07 |
|  | HSA-69278 | Cell Cycle, Mitotic | 0.18 | 1.27E-05 |
|  | HSA-68886 | M Phase | 0.18 | 4.10E-04 |
|  | HSA-72163 | mRNA Splicing - Major Pathway | 0.18 | 4.84E-02 |
|  | HSA-73894 | DNA Repair | 0.17 | 4.20E-03 |
|  | HSA-72203 | Processing of Capped Intron-Containing Pre-mRNA | 0.17 | 1.54E-02 |
|  | HSA-68882 | Mitotic Anaphase | 0.17 | 3.01E-02 |
|  | HSA-68877 | Mitotic Prometaphase | 0.17 | 3.20E-02 |
|  | HSA-72172 | mRNA Splicing | 0.17 | 4.38E-02 |
|  | HSA-199991 | Membrane Trafficking | 0.16 | 1.61E-05 |
|  | HSA-983169 | Class I MHC mediated antigen processing & presentation | 0.16 | 3.50E-03 |
|  | HSA-195258 | RHO GTPase Effectors | 0.16 | 1.16E-02 |
|  | HSA-1852241 | Organelle biogenesis and maintenance | 0.16 | 1.24E-02 |
|  | HSA-69620 | Cell Cycle Checkpoints | 0.16 | 2.27E-02 |
|  | HSA-5653656 | Vesicle-mediated transport | 0.15 | 7.30E-05 |
| MDA-MB-231 | HSA-3322077 | Glycogen synthesis | 1.07 | 2.26E-02 |
|  | HSA-70171 | Glycolysis | 0.82 | 8.70E-05 |
|  | HSA-70326 | Glucose metabolism | 0.76 | 8.70E-05 |
|  | HSA-71387 | Metabolism of carbohydrates | 0.51 | 8.70E-05 |
| HepG2 | HSA-400253 | Circadian Clock | 0.36 | 5.70E-03 |
|  | HSA-8953854 | Metabolism of RNA | 0.17 | 6.93E-05 |
|  | HSA-446203 | Asparagine N-linked glycosylation | 0.17 | 3.65E-02 |
|  | HSA-71387 | Metabolism of carbohydrates | 0.17 | 4.96E-02 |
|  | HSA-199991 | Membrane Trafficking | 0.15 | 8.10E-04 |
|  | HSA-5653656 | Vesicle-mediated transport | 0.15 | 1.00E-03 |
|  | HSA-68886 | M Phase | 0.15 | 4.96E-02 |
|  | HSA-1640170 | Cell Cycle | 0.14 | 1.70E-03 |
|  | HSA-69278 | Cell Cycle, Mitotic | 0.14 | 8.10E-03 |
|  | HSA-74160 | Gene expression (Transcription) | 0.12 | 6.93E-05 |
|  | HSA-597592 | Post-translational protein modification | 0.11 | 8.50E-05 |
|  | HSA-392499 | Metabolism of proteins | 0.1 | 6.93E-05 |
|  | HSA-73857 | RNA Polymerase II Transcription | 0.1 | 1.20E-03 |
|  | HSA-1430728 | Metabolism | 0.09 | 1.50E-04 |
|  | HSA-1643685 | Disease | 0.09 | 4.20E-03 |
|  | HSA-212436 | Generic Transcription Pathway | 0.09 | 2.13E-02 |
| HKC8 | HSA-70263 | Gluconeogenesis | 1.34 | 3.40E-03 |
|  | HSA-70268 | Pyruvate metabolism | 1.28 | 3.84E-02 |
|  | HSA-70171 | Glycolysis | 1.21 | 1.50E-04 |
|  | HSA-71387 | Metabolism of carbohydrates | 0.78 | 7.70E-04 |
| K562 | HSA-9694548 | Maturation of spike protein | 0.64 | 2.65E-02 |
|  | HSA-180746 | Nuclear import of Rev protein | 0.59 | 2.80E-02 |
|  | HSA-168333 | NEP/NS2 Interacts with the Cellular Export Machinery | 0.58 | 4.82E-02 |
|  | HSA-9694635 | Translation of structural proteins | 0.56 | 2.36E-02 |
|  | HSA-70171 | Glycolysis | 0.55 | 2.10E-03 |
|  | HSA-3371556 | Cellular response to heat stress | 0.54 | 4.40E-04 |
|  | HSA-432722 | Golgi Associated Vesicle Biogenesis | 0.53 | 1.17E-02 |
|  | HSA-3371453 | Regulation of HSF1-mediated heat shock response | 0.51 | 7.80E-03 |
|  | HSA-70326 | Glucose metabolism | 0.48 | 4.30E-03 |
|  | HSA-199992 | trans-Golgi Network Vesicle Budding | 0.45 | 3.51E-02 |
|  | HSA-72202 | Transport of Mature Transcript to Cytoplasm | 0.44 | 2.55E-02 |
|  | HSA-159236 | Transport of Mature mRNA derived from an Intron-Containing Transcript | 0.44 | 3.86E-02 |
|  | HSA-72306 | tRNA processing | 0.38 | 3.91E-02 |
|  | HSA-9679506 | SARS-CoV Infections | 0.36 | 1.46E-02 |
|  | HSA-8856828 | Clathrin-mediated endocytosis | 0.35 | 2.57E-02 |
|  | HSA-983231 | Factors involved in megakaryocyte development and platelet production | 0.34 | 2.69E-02 |
|  | HSA-72203 | Processing of Capped Intron-Containing Pre-mRNA | 0.3 | 1.09E-02 |
|  | HSA-2262752 | Cellular responses to stress | 0.24 | 2.70E-03 |
|  | HSA-199991 | Membrane Trafficking | 0.22 | 4.30E-03 |
|  | HSA-8953854 | Metabolism of RNA | 0.21 | 4.30E-03 |
|  | HSA-5653656 | Vesicle-mediated transport | 0.2 | 7.80E-03 |
|  | HSA-597592 | Post-translational protein modification | 0.14 | 8.60E-03 |
|  | HSA-392499 | Metabolism of proteins | 0.11 | 1.09E-02 |
| LNCaP | HSA-72766 | Translation | 0.4 | 2.56E-02 |
| PC3 | HSA-199991 | Membrane Trafficking | 0.11 | 4.90E-03 |
|  | HSA-9675108 | Nervous system development | 0.11 | 1.17E-02 |
|  | HSA-422475 | Axon guidance | 0.11 | 1.28E-02 |
|  | HSA-9006934 | Signaling by Receptor Tyrosine Kinases | 0.11 | 1.73E-02 |
|  | HSA-5653656 | Vesicle-mediated transport | 0.1 | 1.02E-02 |
|  | HSA-1640170 | Cell Cycle | 0.09 | 2.40E-02 |
|  | HSA-597592 | Post-translational protein modification | 0.08 | 1.90E-03 |
|  | HSA-392499 | Metabolism of proteins | 0.07 | 1.20E-03 |
|  | HSA-74160 | Gene expression (Transcription) | 0.07 | 6.50E-03 |
|  | HSA-1643685 | Disease | 0.07 | 1.10E-02 |
|  | HSA-73857 | RNA Polymerase II Transcription | 0.07 | 1.15E-02 |
|  | HSA-212436 | Generic Transcription Pathway | 0.07 | 2.40E-02 |
|  | HSA-1430728 | Metabolism | 0.05 | 1.30E-02 |

This was found by inputting the gathered genes from into STRING and performing a Reactome pathway enrichment. The top Reactome pathways from each cell line were gathered and displayed in the table format here. These pathways were also sorted in terms of their strength. As can be seen, several of the pathways are related to those mediated by HIF1A in cancer cells.

**Supplementary Table S6.** **A ChipSeeker (1) representation of the most enriched pathways in all of the cell lines except HUVEC.**

| Cell Lines | GO terms | Gene ratio | Bg ratio | P-value | Adjusted p-value | q-value |
| --- | --- | --- | --- | --- | --- | --- |
| 501A | RHO GTPase cycle | 263/4647 | 449/10867 | 4.45e-12 | 6.74e-09 | 4.69e-09 |
|  | Antigen processing: Ubiquitination & Proteasome degradation | 189/4647 | 309/10867 | 3.19e-11 | 2.42e-08 | 1.68e-08 |
|  | M Phase | 243/4647 | 418/10867 | 8.54e-11 | 4.31e-08 | 2.99e-08 |
|  | Diseases of signal transduction by growth factor receptors and second messengers | 249/4647 | 433/10867 | 2.22e-10 | 8.41e-08 | 5.85e-08 |
|  | Transcriptional Regulation by TP53 | 214/4647 | 365/10867 | 4.16e-10 | 1.26e-07 | 8.76e-08 |
| MDA-MB-231 | Glucose metabolism | 18/394 | 91/10867 | 3.31E-09 | 1.93e-06 | 1.74e-06 |
|  | Glycolysis | 16/394 | 71/10867 | 3.40E-09 | 1.93e-06 | 1.74e-06 |
|  | Metabolism of carbohydrates | 31/394 | 295/10867 | 9.24E-08 | 3.51e-05 | 3.15e-05 |
|  | Glycogen synthesis | 6/394 | 16/10867 | 1.29E-05 | 3.66e-03 | 3.29e-03 |
|  | Gluconeogenesis | 8/394 | 34/10867 | 2.20E-05 | 5.02e-03 | 4.51e-03 |
| HepG2 | Circadian Clock | 47/3602 | 70/10867 | 5.70E-09 | 8.63E-06 | 6.70E-06 |
|  | DNA Repair | 158/3602 | 335/10867 | 4.58E-08 | 3.46E-05 | 2.69E-05 |
|  | Translation | 135/3602 | 291/10867 | 1.37E-06 | 6.89E-04 | 5.35E-04 |
|  | Asparagine N-linked glycosylation | 137/3602 | 304/10867 | 7.75E-06 | 2.35E-03 | 1.82E-03 |
|  | Chromatin modifying enzymes | 125/3602 | 274/10867 | 9.32E-06 | 2.35E-03 | 1.82E-03 |
| HKC8 | Glycolysis | 11/90 | 71/10867 | 1.15E-11 | 5.27E-09 | 5.14E-09 |
|  | Glucose metabolism | 11/90 | 91/10867 | 1.85E-10 | 3.58E-08 | 3.49E-08 |
|  | Metabolism of carbohydrates | 17/90 | 295/10867 | 2.33E-10 | 3.58E-08 | 3.49E-08 |
|  | Gluconeogenesis | 6/90 | 34/10867 | 3.04E-07 | 3.50E-05 | 3.41E-05 |
|  | Pyruvate metabolism | 4/90 | 31/10867 | 1.17E-04 | 1.07E-02 | 1.05E-02 |
| K562 | Glycolysis | 23/1177 | 71/10867 | 7.98E-07 | 5.66E-04 | 5.16E-04 |
|  | Processing of Capped Intron-Containing Pre-mRNA | 52/1177 | 244/10867 | 1.10E-06 | 5.66E-04 | 5.16E-04 |
|  | Translation of Structural Proteins | 18/1177 | 49/10867 | 1.61E-06 | 5.66E-04 | 5.16E-04 |
|  | mRNA Splicing - Major Pathway | 42/1177 | 183/10867 | 1.61E-06 | 5.66E-04 | 5.16E-04 |
|  | Glucose metabolism | 26/1177 | 91/10867 | 2.33E-06 | 6.58E-04 | 6.00E-04 |
| LNCaP | Chromatin modifying enzymes | 31/578 | 274/10867 | 5.59E-05 | 5.59E-05 | 2.24E-02 |
|  | Chromatin organization | 31/578 | 274/10867 | 5.59E-05 | 5.59E-05 | 2.24E-02 |
|  | Pre-NOTCH Expression and Processing | 17/578 | 109/10867 | 5.81E-05 | 5.81E-05 | 2.24E-02 |
|  | Pre-NOTCH Transcription and Translation | 15/578 | 93/10867 | 1.06E-04 | 1.06E-04 | 3.07E-02 |
| PC3 | RHO GTPase cycle | 403/7682 | 449/10867 | 2.55E-23 | 3.90E-20 | 2.20E-20 |
|  | Diseases of signal transduction by growth factor receptors and second messengers | 377/7682 | 433/10867 | 1.67E-16 | 1.28E-13 | 7.22E-14 |
|  | Antigen processing: Ubiquitination & Proteasome degradation | 276/7682 | 309/10867 | 1.30E-15 | 6.61E-13 | 3.72E-13 |
|  | Transcriptional Regulation by TP53 | 318/7682 | 365/10867 | 3.52E-14 | 1.34E-11 | 7.58E-12 |
|  | Intracellular signaling by second messengers | 271/7682 | 309/10867 | 4.94E-13 | 1.51E-10 | 8.52E-11 |

A pathway analysis was done using ChipSeeker to verify the enriched gene processes.

**Supplementary Table S7. TF cofactors identified between pairs of cell lines.**

| Cell lines | 501A | MDA-MB-231 | HepG2 | HKC8 | HUVEC | K562 | LNCaP | PC3 |
| --- | --- | --- | --- | --- | --- | --- | --- | --- |
| 501A |  | 19/45 = 0.42 | 15/29 = 0.52 | 3/46 = 0.07 | 0/9 = 0.0 | 12/37 = 0.32 | 6/61 = 0.1 | 14/26 = 0.54 |
| MDA-MB-231 | 7/20 = 0.35 |  | 7/29 = 0.24 | 1/46 = 0.02 | 0/9 = 0.0 | 10/37 = 0.27 | 7/61 = 0.11 | 9/26 = 0.35 |
| HepG2 | 15/20 = 0.75 | 25/45 = 0.56 |  | 5/46 = 0.11 | 0/9 = 0.0 | 13/37 = 0.35 | 9/61 = 0.15 | 12/26 = 0.46 |
| HKC8 | 3/20 = 0.15 | 3/45 = 0.07 | 5/29 = 0.17 |  | 6/9 = 0.67 | 7/37 = 0.19 | 34/61 = 0.56 | 3/26 = 0.12 |
| HUVEC | 0/20 = 0.0 | 0/45 = 0.0 | 0/29 = 0.0 | 6/46 = 0.13 |  | 0/37 = 0.0 | 3/61 = 0.05 | 0/26 = 0.0 |
| K562 | 12/20 = 0.6 | 29/45 = 0.64 | 13/29 = 0.45 | 7/46 = 0.15 | 0/9 = 0.0 |  | 9/61 = 0.15 | 18/26 = 0.69 |
| LNCaP | 2/20 = 0.1 | 7/45 = 0.16 | 3/29 = 0.1 | 15/46 = 0.33 | 2/9 = 0.22 | 4/37 = 0.11 |  | 4/26 = 0.15 |
| PC3 | 14/20= 0.7 | 34/45 = 0.76 | 12/29 = 0.41 | 3/46 = 0.07 | 0/9 = 0.0 | 18/37 = 0.49 | 11/61 = 0.18 |  |

The TFs for every pair of cell lines were compared to identify how unique the TFs discovered were. We found that the TFs discovered were largely unique between a pair of cell lines.

**Supplementary Table S8. A list of the 201 TFs that were identified in our study.**

| **Transcription factors (TFs)** | **Cell lines where the TFs were found** |
| --- | --- |
| ABD-B | HKC8 |
| ABI4 | HKC8 |
| ADR1 | LNCaP |
| ARID3A | HKC8,LNCaP |
| ARNT | HepG2,MDA-MB-231 |
| ARR1 | HKC8,HUVEC |
| ASCL2 | K562,MDA-MB-231,PC3 |
| ATF4 | K562 |
| ATF7 | K562 |
| ATHB-5 | HKC8 |
| AZF1 | HKC8,LNCaP |
| B-H2 | HKC8,HUVEC |
| BACH1 | K562,MDA-MB-231,PC3 |
| BATF | K562,PC3 |
| BATF3 | K562 |
| BCD | HKC8,HUVEC |
| BCL6 | K562 |
| BHLHE40 | HepG2 |
| BHLHE41 | HepG2 |
| BRK | LNCaP |
| BTD | HKC8,LNCaP |
| BZIP911 | HKC8 |
| CAD | LNCaP |
| CG11085 | HKC8,HUVEC |
| CG34031 | HKC8 |
| CG42234 | HKC8 |
| CHA4 | HKC8 |
| CREB1 | HKC8,K562 |
| CREB5 | K562 |
| CTCF | 501A,HKC8,HepG2,LNCaP,PC3 |
| DAL80 | HKC8,HUVEC |
| DAL81 | HKC8,LNCaP |
| DFD | HKC8 |
| E2F4 | 501A,HepG2,K562,PC3 |
| E2F6 | 501A,HepG2,K562,MDA-MB-231,PC3 |
| E2F8 | 501A,K562 |
| EBF1 | LNCaP |
| EGR1 | 501A,HepG2,K562,LNCaP,MDA-MB-231,PC3 |
| EHF | 501A,HepG2 |
| EIP74EF | HKC8,LNCaP |
| ELF1 | 501A,HepG2 |
| ELF3 | 501A,HepG2 |
| ELF4 | 501A,HepG2 |
| ELF5 | 501A,HepG2 |
| ERG | PC3 |
| ESR1 | HKC8,LNCaP |
| ETV2 | PC3 |
| ETV4 | PC3 |
| ETV6 | PC3 |
| EWSR1 | LNCaP,MDA-MB-231 |
| EWSR1-FLI1 | LNCaP,MDA-MB-231 |
| FEV | LNCaP |
| FLI1 | LNCaP,MDA-MB-231,PC3 |
| FOS | K562,MDA-MB-231,PC3 |
| FOSL1 | K562,PC3 |
| FOSL2 | K562,MDA-MB-231,PC3 |
| FOXA2 | LNCaP |
| FOXD3 | HKC8 |
| FOXI1 | LNCaP |
| GABPA | HepG2,K562,LNCaP,PC3 |
| GAL4 | LNCaP |
| GATA1 | HepG2,K562 |
| GATA2 | K562 |
| GATA3 | K562 |
| GATA4 | K562 |
| GCM2 | 501A |
| GLIS1 | K562 |
| GLIS2 | 501A,HepG2,MDA-MB-231,PC3 |
| GLIS3 | K562 |
| GLN3 | HKC8,HUVEC |
| GSC | HKC8,HUVEC |
| RELA | HUVEC |
| HAL9 | HKC8,HUVEC |
| HAP5 | HKC8 |
| HAT5 | HKC8,HUVEC |
| HB | LNCaP |
| HES5 | HepG2,MDA-MB-231 |
| HES7 | HepG2,MDA-MB-231 |
| HEY1 | HepG2,MDA-MB-231 |
| HEY2 | MDA-MB-231 |
| HIF1A | HepG2,MDA-MB-231 |
| HINFP | 501A,HepG2,PC3 |
| HKB | HKC8,LNCaP |
| HMX | HKC8,HUVEC |
| HSF1 | HKC8,HUVEC,LNCaP |
| ID1 | LNCaP |
| IME1 | HKC8,LNCaP |
| INO2 | HUVEC |
| INO4 | HUVEC |
| INSM1 | HKC8,HepG2,LNCaP |
| IRF1 | HKC8 |
| IXR1 | HKC8 |
| JDP2 | PC3 |
| JUN | K562,MDA-MB-231,PC3 |
| JUNB | K562,PC3 |
| JUND | K562,PC3 |
| KLF1 | 501A,HepG2,K562,MDA-MB-231,PC3 |
| KLF12 | K562,MDA-MB-231,PC3 |
| KLF13 | HepG2 |
| KLF14 | 501A,HepG2,K562,PC3 |
| KLF16 | 501A,HepG2,K562,MDA-MB-231,PC3 |
| KLF4 | 501A,HepG2,K562,MDA-MB-231,PC3 |
| KLF5 | 501A,HKC8,HepG2,K562,LNCaP,MDA-MB-231,PC3 |
| LYS14 | HKC8,HUVEC |
| MAC1 | HKC8 |
| MAF | MDA-MB-231,PC3 |
| MAFF | K562,PC3 |
| MAFG | K562,MDA-MB-231,PC3 |
| MAFK | K562,MDA-MB-231,PC3 |
| MATALPHA2 | HKC8 |
| MECOM | K562 |
| MIG1 | HKC8,LNCaP |
| MIG2 | HKC8,LNCaP |
| MIG3 | HKC8,LNCaP |
| MSN2 | HKC8,LNCaP |
| MSN4 | HKC8,LNCaP |
| MYF | LNCaP |
| MYOD1 | K562,PC3 |
| MYOG | K562,PC3 |
| MZF1 | 501A,HepG2,K562,MDA-MB-231,PC3 |
| NFAT5 | K562 |
| NFATC1 | K562 |
| NFATC2 | HKC8,K562 |
| NFATC3 | K562 |
| NFE2 | MDA-MB-231,PC3 |
| NFE2L2 | HKC8, K562,MDA-MB-231,PC3 |
| NHLH1 | 501A,HepG2,K562,LNCaP,PC3 |
| NHP10 | HKC8,LNCaP |
| NKX2-3 | K562,MDA-MB-231 |
| NR1H2 | HKC8 |
| NRF1 | 501A,HepG2,K562,PC3 |
| NUB | HKC8 |
| OC | HKC8,HUVEC |
| ONECUT | HKC8,LNCaP |
| OPA | HKC8,LNCaP |
| PAN | HKC8,LNCaP |
| PAX5 | K562 |
| PBX1 | HKC8 |
| PDR3 | HKC8 |
| PLAG1 | HKC8,HepG2,LNCaP |
| POU5F1 | HKC8 |
| PPARG | HKC8,K562 |
| PRDM1 | K562 |
| PRRX2 | HKC8,HUVEC |
| PTX1 | HKC8,HUVEC |
| MAX | HKC8 |
| MYC | HKC8 |
| REST | HKC8,HUVEC |
| RGM1 | HKC8,LNCaP |
| RME1 | LNCaP |
| RPN4 | HKC8,LNCaP |
| RREB1 | K562 |
| RSC30 | HKC8,LNCaP |
| RUNX1 | MDA-MB-231 |
| RXRA | HKC8,K562 |
| SOX3 | K562 |
| SP1 | 501A,HKC8,HepG2,K562,LNCaP,MDA-MB-231,PC3 |
| SP2 | 501A,HepG2,K562,MDA-MB-231,PC3 |
| SP3 | 501A,HepG2,MDA-MB-231,PC3 |
| SP4 | 501A,HepG2,K562,MDA-MB-231,PC3 |
| SPI1 | HKC8,LNCaP,PC3 |
| SPIB | HKC8,LNCaP,PC3 |
| SPIC | K562,PC3 |
| SPT23 | HKC8,HUVEC,LNCaP |
| SREBF1 | HepG2 |
| STAT1 | HKC8,HUVEC,K562,LNCaP |
| STAT3 | K562,LNCaP,PC3 |
| STAT4 | K562 |
| STAT5A | K562 |
| STB3 | LNCaP |
| STP1 | LNCaP |
| SUT1 | HKC8,LNCaP |
| SWI5 | HKC8,LNCaP |
| TAL1 | HKC8,HepG2 |
| TCF12 | K562,PC3 |
| TCF3 | HKC8 |
| TEAD1 | K562,LNCaP |
| TEAD3 | K562 |
| TFAP2A | 501A,HKC8,HepG2,K562,LNCaP,MDA-MB-231,PC3 |
| TFAP2B | 501A,HepG2,K562,PC3 |
| TFAP2C | 501A,HepG2,K562,PC3 |
| TFEB | HepG2 |
| TFEC | HepG2 |
| TIN | HKC8,HUVEC |
| TP53 | HKC8 |
| UGA3 | HKC8,LNCaP |
| UME6 | LNCaP |
| UNC-4 | HKC8,HUVEC |
| VND | HKC8,HUVEC |
| YKL222C | HKC8 |
| YLL054C | HKC8 |
| YPR022C | LNCaP |
| YPR196W | LNCaP |
| ZBTB7A | 501A,K562 |
| ZBTB7C | 501A,HepG2,K562 |
| ZFX | 501A,HepG2,K562,MDA-MB-231,PC3 |
| ZIC1 | 501A,HepG2,K562,MDA-MB-231,PC3 |
| ZIC3 | 501A,HepG2,K562,MDA-MB-231,PC3 |
| ZIC4 | 501A,HepG2,K562,MDA-MB-231,PC3 |
| ZNF263 | 501A,HepG2,K562,MDA-MB-231,PC3 |
| ZNF740 | 501A,HepG2,K562,MDA-MB-231,PC3 |

Here, we compiled the list of all TF cofactor with their motifs similar to a predicted motif with a STAMP E-value < 1E-5 and labeled which cell line(s)the cofactors were found in. Because of the strict STAMP E-value cutoff and the degenerate nature of motifs, a cofactor is likely to be active in cell lines not listed in the second column as well.

**Supplementary Table S9. A list of other high potential TFs that may function as cofactors.**

| Transcription factors | Supporting evidence |
| --- | --- |
| NRF2 (NFE2L2) | Directly supported by external evidence ^1^  Common target genes ^2^ |
| KLF5 | Directly supported by external evidence ^3^ |
| TFAP2A | Directly supported by external evidence ^4^ |
| EGR1 | Directly supported by external evidence ^5^ |
| ZNF263 | Likely: Discovered to have a significantly enriched motif near HIF1A binding sites in HKC8 and RCC4 cells ^6^ |
| ZNF740 | Paralog of ZNF197, a known TF cofactor as found in BioGRID ^7^ |
| E2F6 | Reasons mentioned in main text – common target genes, integral to HIF1A’s repression functions |
| E2F4 | Reasons mentioned in main text – common target genes, integral to HIF1A’s repression functions |
| HSF1 | Very likely: Shows high potential due to the co-regulation between the two TFs ^8^ |
| STAT1 | Supported by the PIP database (11), and is interconnected with HIF1A’s regulatory functions |
| STAT5A | Very likely: STAT5A is closely related to tumorigenesis ^9^, upregulates EPAS1, a known HIF1A TF cofactor^7^, is itself upregulated by HIF1A ^10^, found in the PIP database ^11^ |
| FOSL2 | Discovered to have a significantly enriched motif near HIF1A binding sites using ENCODE data ^6^ |
| GABPA | Likely: AMP kinase coordinates tumor bioenergetics through the regulation of GABPA and HIF1A ^12^ |
| NRF1 | Very likely for reasons mentioned in study ^13^ |
| JUND | Directly supported by external evidence ^14^ |
| FLI1 and EWSR1-FLI1 | Directly supported by external evidence ^15^  In the data gathered, the chimeric protein EWSR1-FLI1 was also found. Since EWSR1-FLI1 has regulatory functions that are essential to tumorigenesis ^16^, FLI may interact with HIF1A individually or through the EWSR1-FLI1 TF complex. |
| TEAD1 | Directly supported by external evidence: Shown to form a TF-complex with YAP and HIF1A ^17^ |
| GATA2 | Forms a complex with AP-1 and HIF1A ^18^ |
| HEY1 | Directly supported by external evidence ^19,20^  TF complex plays essential role in Mitochondrial genesis |
| FOXA2 | Directly supported by external evidence ^21^ |
| IRF1 | Directly supported by external evidence ^22^ |
| ETV4 | Directly supported by external evidence ^23^  ETV4 co-regulates many of HIF1A’s target genes |
| PPARG | Directly supported by external evidence ^24^ |

The top TFs that were not found in any of the large PPI databases were organized in a table format with supporting evidence of their role as a potential TF cofactor.
